# Piperlongumine (PL) conjugates induce targeted protein degradation

**DOI:** 10.1101/2022.01.21.474712

**Authors:** Jing Pei, Yufeng Xiao, Xingui Liu, Wanyi Hu, Amin Sobh, Yaxia Yuan, Shuo Zhou, Nan Hua, Samuel G Mackintosh, Xuan Zhang, Kari B. Basso, Manasi Kamat, Qingping Yang, Jonathan D. Licht, Guangrong Zheng, Daohong Zhou, Dongwen Lv

**Author notes:** **Correspondence: Email address(es) of corresponding author(s)** Guangrong Zheng, Daohong Zhou, Dongwen Lv. These authors contributed equally (Jing Pei, Yufeng Xiao, Xingui Liu). **LEAD CONTACT:** Dongwen Lv.

## Abstract

PROteolysis Targeting Chimeras (PROTACs) are bifunctional molecules that degrade target proteins through recruiting E3 ligases. However, their application is limited in part because few E3 ligases can be recruited by known E3 ligase ligands. In this study, we identified piperlongumine (PL), a natural product, as a covalent E3 ligase recruiter, which induces CDK9 degradation when it is conjugated with SNS-032, a CDK9 inhibitor. The lead conjugate **955** can potently degrade CDK9 in a ubiquitin-proteasome-dependent manner and is much more potent than SNS-032 against various tumor cells *in vitro*. Mechanistically, we identified KEAP1 as the E3 ligase recruited by **955** to degrade CDK9 through a TurboID-based proteomics study, which was further confirmed by KEAP1 knockout and the nanoBRET ternary complex formation assay. In addition, PL-Ceritinib conjugate can degrade EML4-ALK fusion oncoprotein, suggesting that PL may have a broader application as a covalent E3 ligase ligand in targeted protein degradation.

## INTRODUCTION

PROteolysis TArgeting Chimeras (PROTACs) are potentially more potent anticancer therapeutics than small molecule inhibitors (SMIs) because they can degrade oncoproteins in an event-driven manner.^1,2^ Moreover, compared to SMIs that only block the catalytic function of proteins of interest (POIs), PROTACs can further remove the scaffold function of POIs by inducing their degradation. Furthermore, PROTACs can target some previously considered undruggable proteins, such as transcription factors. For example, a potent signal transducer and activator of transcription 3 (STAT3) PROTAC has been generated and shown efficacy *in vivo.*^3^ In addition, PROTAC-induced POI degradation is driven by the ternary complex formation and can be affected by the availability of lysine on the POI.^4–6^ Therefore, PROTACs can be more specific/selective than their SMI predecessors. Because of these advantages, to date, more than 10 PROTACs have been advanced to phase I or phase II clinical trials.^7^ The targets include androgen receptor (AR), estrogen receptor (ER), B-cell lymphoma extra-large (BCL-xL), bruton tyrosine kinase (BTK), bromodomain-containing protein 9 (BRD9), interleukin-1 receptor-associated kinase 4 (IRAK4), STAT3, and tropomyosin receptor kinase (TRK).^7^

Despite great progress in the field, there are still some obstacles that prevent PROTACs from being more useful.^8^ Among them, to date, only a few E3 ligases and ligands are available to generate PROTACs. The human genome encodes more than 600 E3 ligases,^9^ and only a few of them (CRBN,^2^ VHL,^1^ cIAPs,^10^ and MDM2^11^) have been utilized by PROTACs to degrade POIs. This limits the ability to generate PROTACs for a POI that is not a suitable neo-substrate for those E3 ligases because different proteins may require different E3 ligases to effectively mediate their degradation. For example, endogenous KRAS^G12C^ can be degraded by VHL-recruiting PROTACs^12^ rather than CRBN-recruiting PROTACs.^13^ In addition, some E3 ligases are highly expressed in certain tumor cells,^13^ which may provide the opportunity to selectively degrade POIs in those tumor cells if ligands of those E3 ligases can be found. Recent studies have also shown that cancer cells develop resistance to VHL-based bromodomain and extra-terminal domain (BET) PROTACs due to loss of CUL2, as well as to CRBN-based BET and CDK9 PROTACs because of CRBN loss.^14,15^ Therefore, significant efforts have been devoted to identify new E3 ligase ligands, resulting in the discovery of a number of new E3 ligase ligands that can recruit AhR,^16^ DCAF11,^17^ DCAF15,^18^ DCAF16,^19^ FEM1B,^20^ KEAP1,^21,22^ RNF114,^23,24^ and RNF4^25^ E3 ligases to degrade POIs. Based on mathematical modeling^26^ and previous studies,^19,21,23,27,28^ covalent E3 ligase ligand-based PROTACs may outperform non-covalent E3 ligase ligand-based PROTACs due to better kinetics of ternary complex formation, and minimal perturbation of its endogenous substrates with low fractional occupancy of the recruited E3 ligase. In addition, accumulating evidence shows that a covalent E3 ligase ligand in a PROTAC provides additional selectivity for a given POI^19,23,25^. Identifying more E3 ligase ligands can further expand the toolbox, overcome the drug resistance, and potentially generate more potent and specific PROTACs.

Piperlongumine (PL, Figure S1A) is a natural product that exhibits potent antitumor activity,^29^ in part via induction of oxidative stress through its two Michael acceptors that can covalently react with GSTP1^30^ and GSTO1.^31^ Our previous studies also showed that PL can selectively kill senescent cells in part through induction of OXR1 degradation in a proteasome-dependent manner.^32,33^ In addition, we found that PL can bind several intracellular proteins in senescent cells, including 8 different E3 ligases,^32^ suggesting that PL could be used to recruit E3 ligase and generate PROTACs. SNS-032 is a CDK inhibitor with antitumor activity. It can potently inhibit CDK9 (IC_50_ = 4 nM), CDK2 (IC_50_ = 38 nM), and CDK7 (IC_50_ = 62 nM), and also has moderate inhibitory activity for a panel of other kinases.^34^ Interestingly, when it was conjugated with thalidomide, the conjugate (THAL-SNS-032) became more specific and could only degrade CDK9 but not other known SNS-032 targets.^35^ To test if PL can be used as a new covalent E3 ligase ligand, we linked PL to SNS-032, and found that the PL-SNS-032 conjugates potently induced CDK9 degradation. Mechanistically, we demonstrated that the lead PL-SNS-032 conjugate, **955**, degraded CDK9 in a ubiquitin-proteasome system (UPS) dependent manner. Using the TurboID-based proteomics combined with siRNA knockdown, CRISPR-Cas9 gene knockout, and the nanoBRET ternary complex formation assay, we identified and confirmed KEAP1 as the E3 ligase recruited by **955** to degrade CDK9. Furthermore, a PL-based ALK PROTAC can potently degrade ALK-fusion protein in ALK positive non-small cell lung cancer (NSCLC) cells. Our results demonstrate that PL has the potential to be used as a new covalent E3 ligase ligand to generate PROTACs to degrade different target proteins through recruiting KEAP1. In addition, our study reveals that the TurboID-based proteomics is a very useful technology to identify E3 ligases recruited by natural products for targeted protein degradation.

## RESULTS AND DISCUSSION

### Generation of PL-conjugated CDK9 PROTACs

Our previous study showed that PL can pull down 8 E3 ligases in senescent cells.^32^ Here, we used a competitive activity-based protein profiling (ABPP) assay with the PL-Alkyne as a probe to validate whether PL can covalently bind E3 ligases in cancer cells as well (Figure S1A). The assay was done using MOLT4 human T-cell acute lymphoblastic leukemia (T-ALL) cells as illustrated in Figure S1B and described in STAR Methods. To exclude proteins that bound non-specifically to PL-Alkyne, we also included a sample in which the cells were pre-treated with a high concentration of PL before the addition of PL-Alkyne to compete for protein binding (Figure S1B). The western blot result showed that PL-Alkyne pulled down many proteins, which could be effectively competed by the pre-treatment with PL (Figure S1C), suggesting that most of the pulled-down proteins by the probe are PL-binding proteins. The mass spectrometry (MS) results showed that PL can bind to about 300 proteins (Figure S1D and Table S1), including GSTO1, a previously identified PL target.^31^ We performed gene ontology (GO) and Kyoto encyclopedia of genes and genomes (KEGG) pathway enrichment analyses of those PL-binding proteins and found that many of the proteins involved in the UPS were highly enriched (Figure S1E-H and Table S1), including 9 E3 ligases (Figure S1D). These findings indicate that PL has the potential to be used as a new E3 ligase ligand for PROTAC design.

CDK9 is a well-established cancer target and can be effectively degraded by a PROTAC (THAL-SNS-032) consisting of thalidomide, a linker, and the CDK inhibitor SNS-032.^35^ To test our hypothesis, we generated a series of PL-SNS-032 bifunctional molecules with linkers of different types and lengths (Figure S2A, B) and examined their potencies in degrading CDK9 in MOLT4 cells (Figure S2A-C). These preliminary assays identified **955** as a lead compound for further evaluation and characterization because of its high potency in these assays (Figure 1A). Especially, **955** potently degraded CDK9 with a DC_50_ value of 9 nM after 16 h treatment in MOLT4 cells (Figure S2B, C). Even with a short-term treatment (6 h), **955** was able to potently degrade CDK9 while the warhead SNS-032 could not (Figure 1B). In addition, **955** was more potent in inducing PARP cleavage than SNS-032 (Figure 1B). The time-course study showed that **955** induced almost complete degradation of CDK9 at 0.1 µM and within 8 h of treatment (Figure 1C) and its effect lasted up to 18 h after the removal of **955** from the culture (Figure 1D). Similar results were also observed in 293T cells (Figure S3A) and K562 cells (Figure S3B). To exclude the possibility that the degradation of CDK9 is caused by the combination effect of PL and SNS-032, we treated MOLT4 cells with either PL or SNS-032 alone or their combination and found that none of these treatments degraded CDK9 (Figure 1E). However, pre-treatment of MOLT4 cells with PL or SNS-032 blocked **955**-induced CDK9 degradation (Figure 1F).

**Figure 1.**
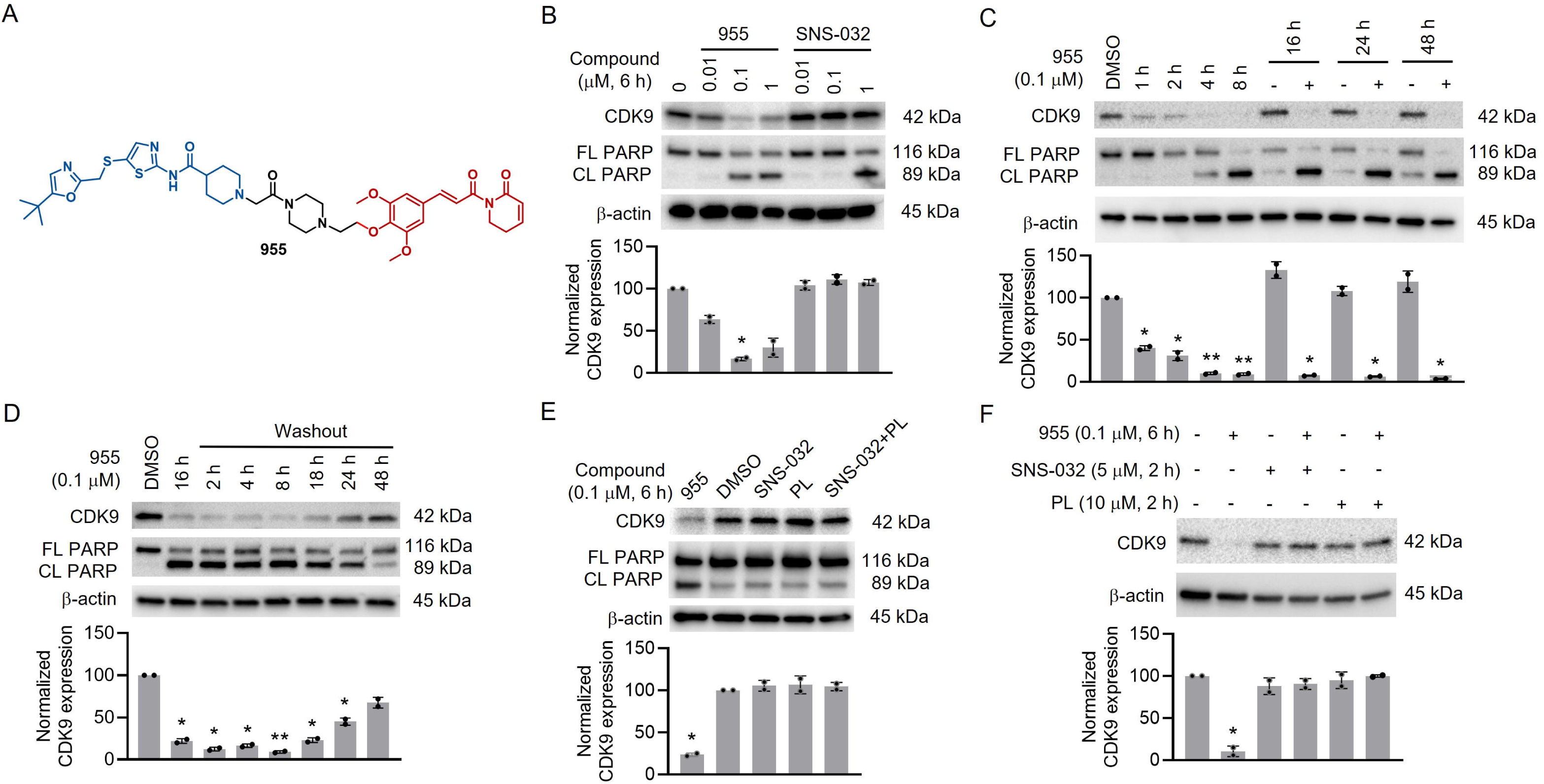
The lead PL-SNS-032 conjugate 955 potently induces CDK9 degradation and apoptosis. (A) The structure of **955**; (B) **955** but not SNS-032 induces the degradation of CDK9 and is more potent than SNS-032 in the induction of PARP cleavage; (C) Time course of **955**-induced CDK9 degradation and PARP cleavage; (D) **955** induces CDK9 degradation and PARP cleavage for up to 18 h after washout of **955**; (E) PL alone or in combination with SNS-032 does not induce CDK9 degradation; and (F) Pre-treatment with SNS-032 or PL blocks the CDK9 degradation induced by **955**. All the experiments were performed in MOLT4 cells. Representative immunoblots are shown and β-actin was used as an equal loading control in all immunoblot analyses. FL PARP (full-length PARP) and CL PARP (cleaved PARP) are used as indicators of apoptosis. The quantification of the relative CDK9 protein content in the immunoblots is presented as mean ± SD (*n* = 2 biologically independent experiments) in the bar graph (bottom panel). Statistical significance was calculated with unpaired two-tailed Student’s *t*-tests. \**P* < 0.05; \*\**P* < 0.01. See also Figures S2 and S3.

### Mechanism of action of 955 in degrading CDK9

To confirm that **955** functions as a PROTAC to degrade CDK9, we performed a series of mechanistic studies using inhibitors to block different protein degradation pathways. First, we treated MOLT4 cells with vehicle, the proteasome inhibitor MG132, or bortezomib prior to **955** treatment. The results showed that the degradation of CDK9 can be blocked by the two different proteasome inhibitors (Figure 2A). In contrast, pre-treatment with either of the two lysosome inhibitors, Baf-A1 and chloroquine, or with the pan-caspase inhibitor QVD does not block CDK9 degradation by **955** (Figure 2B, C). These results demonstrate that **955** induces CDK9 degradation through the proteasome but not lysosome and activated caspases. Furthermore, we used two E1 inhibitors, TAK-243 and PYR-41, to verify that **955**-induced CDK9 degradation is E1-dependent (Figure 2D, E). In our competitive ABPP assay, 9 PL-recruited E3 ligases were identified by the LC-MS/MS (Figure S1D), and thus we next investigated which type of E3 ligase(s) is involved in the degradation of CDK9 by **955**. MLN4924 is a NEDD8-activating enzyme (NAE) inhibitor and can selectively inhibit cullin RING-related ubiquitin E3 ligase(s) (CRLs) through blocking the neddylation of cullin.^36,37^ We found that MLN4924 pre-treatment blocks **955**-induced CDK9 degradation (Figure 2F), suggesting that a CRL is likely to be recruited by **955** to degrade CDK9. Similar results were also observed in 293T and K562 cells (Figure S3C-F). Collectively, these findings suggest that **955** degrades CDK9 in a UPS-dependent manner probably via recruiting a CRL. Furthermore, we investigated whether **955** degrading CDK9 relies on PL to covalently recruit an E3 ligase via its two Michael acceptors by generating **336**, in which the C2-C3 and C7-C8 double bonds of PL are reduced (Figure 2G). We found that **955**, but not **336,** can induce CDK9 degradation (Figure 2H), indicating that **955** degrades CDK9 via covalently recruiting a CRL through PL’s Michael acceptors.

**Figure 2.**
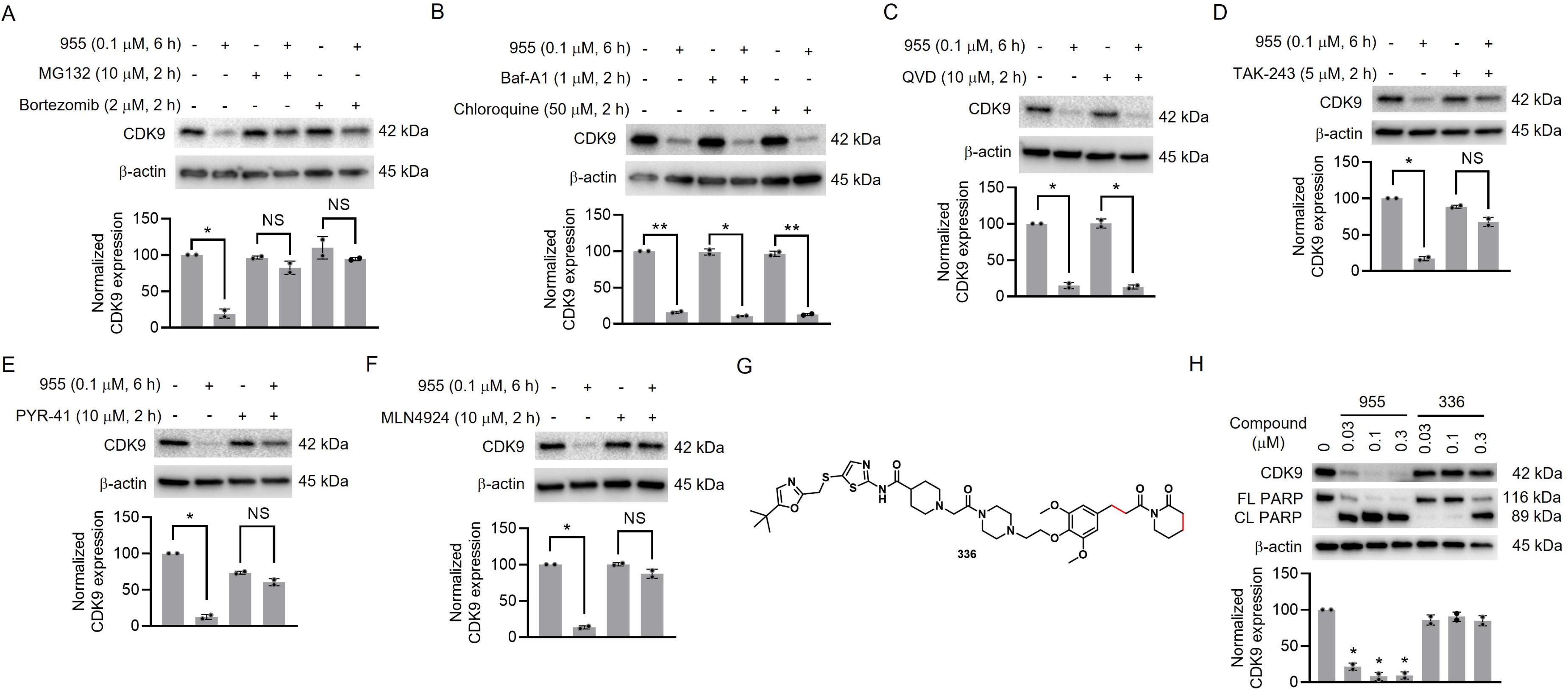
Mechanism of action of 955 in degrading CDK9. (A) Pre-treatment with the proteasome inhibitor MG132 or bortezomib blocks CDK9 degradation by **955**; (B) Pre-treatment with the lysosome inhibitor Baf-A1 or chloroquine does not block CDK9 degradation by **955**; (C) Pre-treatment with pan-caspase inhibitor QVD does not block CDK9 degradation by **955**; (D-E) Pre-treatment with E1 inhibitor TAK-243 (D) or PYR-41 (E) blocks CDK9 degradation by **955**; (F) Pre-treatment with neddylation inhibitor MLN4924 blocks CDK9 degradation by **955**. (G) The structure of **336** illustrates the reduction of the C2-C3 and C7-C8 double bonds of PL in comparison with **955**; (H) **955,** but not **336**, can degrade CDK9. All the experiments were performed in MOLT4 cells. Representative immunoblots are shown and β-actin was used as a loading control in all immunoblot analyses. The quantification of the relative CDK9 protein content in the immunoblots is presented as mean ± SD (*n* = 2 biologically independent experiments) in the bar graph (bottom panel). Statistical significance was calculated with unpaired two-tailed Student’s *t*-tests. \**P* < 0.05; \*\**P* < 0.01; NS: not significant. See also Figure S3.

### Identification of the E3 ligase(s) recruited by 955 to degrade CDK9

PROTACs induce the degradation of target proteins by hijacking an E3 ligase. However, it is usually difficult to directly identify the E3 ligase recruited by a new E3 ligase ligand in a PROTAC using the co-immunoprecipitation method because the formation of the complex is transient and very dynamic.^38^ TurboID assay is a powerful proximity labeling method that can sensitively detect weak and transient protein-protein interactions by biotinylating proteins that interact with a bait protein fused with an engineered biotin ligase.^39^ Similar biotin-based proximity labeling assays, such as APEX2 assay^40^ and AirID assay,^41^ have been used to study the degrader-induced or inhibitor-blocked E3 ligase:substrate interactions in cells. Therefore, we adapted TurboID technology to identify and characterize the specific E3 ligase(s) recruited by **955** to mediate CDK9 degradation by ectopically expressing V5-TurboID-CDK9 in 293T cells and comparing the biotinylated proteins with or without **955** treatment (Figure 3A). The results from this assay showed that **955** can induce the interaction of CDK9 with 6 different E3 ligases (Figure 3B, Table S2). To compare the PL-binding E3 ligases identified by the competitive ABPP assay (Figure 1D) with those identified by the TurboID-bait assay (Figure 3B), we found that KEAP1 is the only CRL identified by both methods. A previous study showed that KEAP1 can be recruited to degrade Tau protein by a KEAP1-based peptide PROTAC.^42^ In addition, two recent studies discovered KEAP1 E3 ligase ligands that can be used to recruit KEAP1 to degrade BRD4 after linking to the BRD4 inhibitor JQ1.^21,22^ To determine whether KEAP1 is the E3 ligase recruited by **955** to degrade CDK9, we first used immunoblot to confirm the results of the TurboID-bait assay. We found that significantly more KEAP1 can be pulled down after **955** treatment than without **955** treatment in the presence of biotin. By contrast, β-actin cannot be pulled down with or without 955 treatment under the same experimental conditions (Figure 3C). To further validate that KEAP1 is recruited by **955** to mediate CDK9 degradation, we treated MOLT4 cells with the known KEAP1 covalent inhibitors CDDO-ME, CDDO-IM, and dimethyl fumarate (DMF),^43^ prior to the addition of **955**. Treatment with these inhibitors completely blocked **955**-induced CDK9 degradation (Figure 3D, E). Furthermore, we used both siRNA and CRISPR-Cas9 to knock down/out of KEAP1 and found that CDK9 degradation induced by **955** was blocked after depleting KEAP1 (Figure 3F and S4A). We also knocked down two additional E3 ligases (TRIP12 and TRAF6) identified by TurboID-bait assay using siRNAs, which had no effect on **955**-induced CDK9 degradation (Figure S4B, C). Ectopic expression of Halo-tagged KEAP1 in the KEAP1 stable knockout cells confirmed that the expression of KEAP1 rescued CDK9 degradation induced by **955** (Figure 3G), suggesting KEAP1 is likely the primary E3 ligase recruited by **955** to degrade CDK9. Depleting KEAP1 can stabilize NRF2 as expected (Figure S4A). To rule out the possibility that the upregulation of NRF2 and its downstream pathways are involved in metabolizing **955** to prevent it from being an effective PROTAC degrader, we simultaneously knocked down both KEAP1 and NRF2 in cells. The results from this study show that knocking down NRF2 does not reverse the effect of KEAP1 knocking down on abrogation of **955**-induced CDK9 degradation, confirming that **955** induces CDK9 degradation by recruiting KEAP1 but not by activation of NRF2 (Figure S4D). To further confirm that **955** recruits KEAP1 to degrade CDK9 through covalent binding, we performed a gel-based competitive ABPP assay, in which we used serially diluted **955** to compete with iodoacetamide (IA)-alkyne, a non-selective cysteine-reactive probe, for binding to purified KEAP1 protein. The result from this assay confirms that **955** can compete with IA-alkyne to covalently bind to KEAP1 in a concentration-dependent manner (Figure 3H). Lastly, forming a POI: PROTAC: E3 ternary complex is a necessary step for the induction of POI ubiquitination and degradation. Therefore, we fused CDK9 with HiBit-tag and KEAP1 with Halo-tag for nanoBRET assay to further test whether **955** could induce the formation of the CDK9:**955**:KEAP1 ternary complex in live cells (Figure 3I). The results demonstrated that **955** can induce the formation of the CDK9:**955**:KEAP1 ternary complex in a dose-dependent manner while the induction is negligible by **336**, suggesting that the formation of this ternary complex is dependent on the covalent binding of **955** to KEAP1 via the PL Michael acceptor(s) (Figure 3I). In conclusion, **955** can covalently hijack KEAP1 to mediate CDK9 degradation.

**Figure 3.**
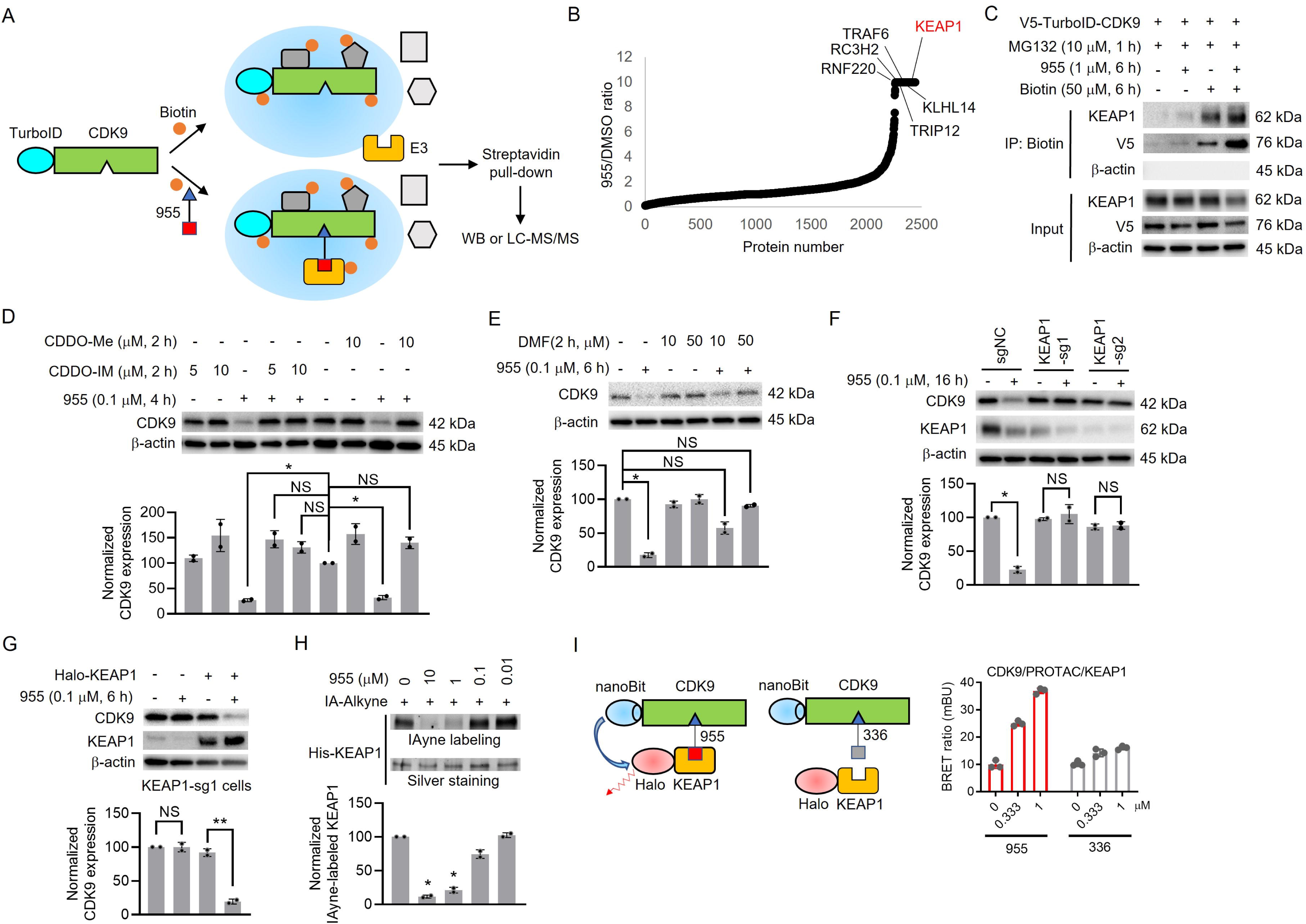
Identification of CDK9 degrading E3 ligase(s) recruited by 955. (A) Schematic illustration of the TurboID-bait assay to identify CDK9 degrading E3 ligase(s) recruited by **955**; (B) The biotinylated proteins identified by MS in 293T cells. E3 ligases are labelled with KEAP1 highlighted in red; (C) Western blot analysis to validate KEAP1 recruitment by **955**. Representative immunoblots from two independent experiments are shown. (D) KEAP1 inhibition with CDDO-ME and CDDO-IM blocks **955**-induced CDK9 degradation in MOLT4 cells; (E) Inhibition of KEAP1 with DMF blocks **955**-induced CDK9 degradation in MOLT4 cells. (F) KEAP1 knockout blocks **955**-induced CDK9 degradation. KEAP1 depleted (sg1 and sg2) and control (sgNC) H1299 cells were either untreated or treated with **955** for 16 h. (G) Ectopic expression of Halo-tagged KEAP1 in KEAP1 stable knockout cells rescues **955**-induced CDK9 degradation. KEAP1-sg1 stable knockout H1299 cells were transfected with or without Halo-KEAP1 for 48 h and then the cells were reseeded and treated with 955 for 6 h. Representative immunoblots are shown and β-actin was used as a loading control in all immunoblot analyses. In panels D-G, representative immunoblots are shown and β-actin was used as an equal loading control. The quantification of the relative CDK9 protein content in the immunoblots is presented as mean ± SD (*n* = 2 biologically independent experiments) in the bar graph (bottom panel). Statistical significance was calculated with unpaired two-tailed Student’s *t*-tests. \**P* < 0.05; \*\**P* < 0.01; NS: not significant. (H) Gel-based ABPP assay demonstrates that **955** competes for IA-Alkyne binding to KEAP1 in a dose-dependent manner. Representative immunoblots from two independent experiments are shown. The quantification of the relative IAyne-labeled His-KEAP1 protein content is presented as mean ± SD (*n* = 2) in the bar graph (bottom panel). Statistical significance was calculated with unpaired two-tailed Student’s *t*-tests. \**P* < 0.05. (I) nanoBRET assay demonstrates that **955,** but not **336**, can induce the formation of the intracellular ternary complexes in live 293T cells. Data are expressed as mean ± SD of three biological replicates. See also Figure S4 and Table S2.

### 955 is a more potent anti-cancer agent than SNS-032

To evaluate the anti-cancer efficacy and specificity of **955**, a series of experiments were conducted to compare **955** with its warhead SNS-032. We first compared the global proteome changes induced by **955** and SNS-032 in MOLT4 cells. As expected, cells exhibited a significant reduction of CDK9 after treatment with 0.1 µM **955** for 1 h and 6 h while treatment with 1 µM SNS-032 for 6 h had no significant effect on CDK9 (Figure 4A-C and Table S3). Interestingly, **955**, but not SNS-032, can also potently degrade CDK10 (Figure 4A-D, and Table S3) in a proteasome- and CRL-dependent manner because pre-treatment with MG132 and MLN4924 can block the degradation (Figure 4E). The previously reported CRBN-based CDK9 PROTAC (THAL-SNS-032) also has a similar but weaker effect on inducing CDK10 degradation,^35^ which can be explained by the moderate binding between SNS-032 and CDK10 confirmed by Kinativ screening in MOLT4 lysates.^35^ Because CDK9 plays an important role in regulation of gene transcription, inhibition/degradation of CDK9 can lead to reduction of many short half-life proteins including c-Myc and MCL-1.^44^ These downstream proteins can further regulate the expression of their downstream targets. Therefore, it is not uncommon to see that more proteins are downregulated in cells after longer treatment with a CDK9 inhibitor/degrader than shorter treatment. This may explain why more proteins were downregulated in cells after 6 h treatment with **955** than in cells after 1 h treatment (Figure 4A, B). Importantly, **955** reduced the expression of CDK9 and CDK10 at both time points whereas SNS-032 had no effect on their expression, suggesting that CDK9 and CDK10 are likely the primary and direct targets of **955**. This suggestion is further supported by the findings that most of other proteins were similarly downregulated in cells 6 h after **955** and SNS-032 treatment (Blue dots shown in Figure 4A-C and S5A). Collectively, these findings suggest that **955** can also downregulate CDK9 downstream targets as SNS-032 but via a different mechanism, i.e. via degradation of CDK9.

**Figure 4.**
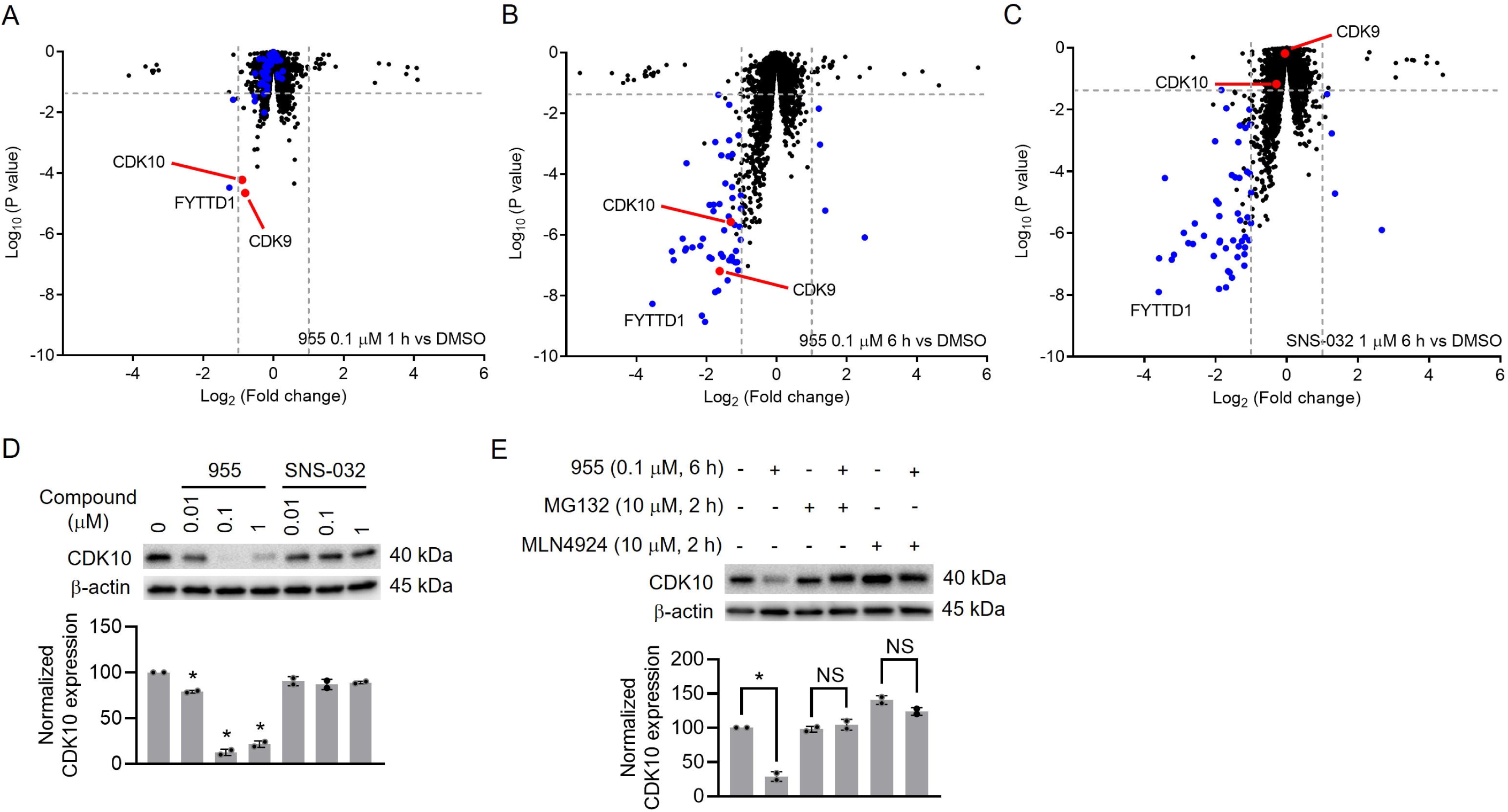
955 selectively degrades CDK9 and CDK10 by proteomic studies. **(A-C)** Proteome changes induced by **955** or SNS-032 in MOLT4 cells; Blue dot: shared differentially expressed proteins after 0.1 μM 955 or 1 μM SNS032 treatment for 6 h. (D) **955**, but not SNS-032, degrades CDK10 in MOLT4 cells. (E) Pre-treatment with MG132 and MLN4924 blocks CDK10 degradation induced by **955**. In panels D and E, representative immunoblots are shown and β-actin was used as an equal loading control. The quantification of the relative CDK10 protein content in the immunoblots is presented as mean ± SD (*n* = 2 biologically independent experiments) in the bar graph (bottom panel). Statistical significance was calculated with unpaired two-tailed Student’s *t*-tests. \**P* < 0.05; NS: not significant. See also Figure S5 and Table S3.

CDK9 plays a key role in regulating gene transcription by interacting with RNA polymerase II.^44^ Therefore, we further performed RNA-seq to analyze the transcriptional changes in the same samples used for the proteome profiling. The results from this analysis confirmed that **955** is more potent than SNS-032 in the down-regulation of CDK9-dependent transcription in MOLT4 cells after 6 h treatment (Figure 5A, B, Figure S5B, C, and Table S4). In addition, MOLT4 cells treated with 0.1 µM **955** and 1 µM SNS-032 displayed very similar proteomic and transcriptomic profiles, suggesting that **955** is more potent than SNS-032 in downregulating CDK9 downstream targets and its effects are not caused by PL. This suggestion is further supported by the analyses of c-Myc and MCL-1, two well-known downstream targets of CDK9.^44^ Specifically, quantitative real-time PCR (qPCR) and immunoblot revealed that **955** is about 10-fold more potent than SNS-032 in downregulation of c-Myc and MCL-1 expression at the levels of both mRNA and protein (Figure 5C, D). We further compared the antiproliferative effect of **955** with SNS-032 in prostate cancer cell lines since previous studies showed that targeting CDK9 is an effective way to inhibit prostate cancer.^44^ Two AR-positive (LNCaP and 22RV1) and two AR-negative (PC3 and DU145) cell lines were studied. We found that **955** exhibits antitumor activities against all four cell lines with EC_50_ values in the single digital nM range, while the EC_50_ values of SNS-032 against these cell lines are over 100 nM (Figure 5E). In addition, PL itself had no anti-proliferative effect within 3 μM and combined treatment of PL and SNS-032 showed a similar EC_50_ to SNS-032 in all the tested prostate cancer cell lines, which further confirmed that **955** is more potent than SNS-032 against the tumor cells (Figure 5E). Furthermore, immunoblot assays showed that **955** potently degraded CDK9 and downregulated c-Myc and MCL-1 in LNCaP cells (Figure 5F). Collectively, these results indicate that **955** is more potent than SNS-032 against tumor cells. To further test if the E3 ligase KEAP1 recruited by **955** is relevant to its antiproliferative activity, we knocked down KEAP1 in LNCaP prostate cancer cells and H1299 lung cancer cells and measured the cell-killing effects in KEAP1 siRNA- and control siRNA-treated cells. The results confirmed that knockdown of KEAP1 attenuated the cytotoxicity of **955**, but had no significant effect of the warhead SNS-032 and compound **336** that does not recruit KEAP1 (Figure 3I and Figure S6). These results provide additional support that **955** kills tumor cells primarily via degrading CDK9 by recruiting the E3 ligase KEAP1.

**Figure 5.**
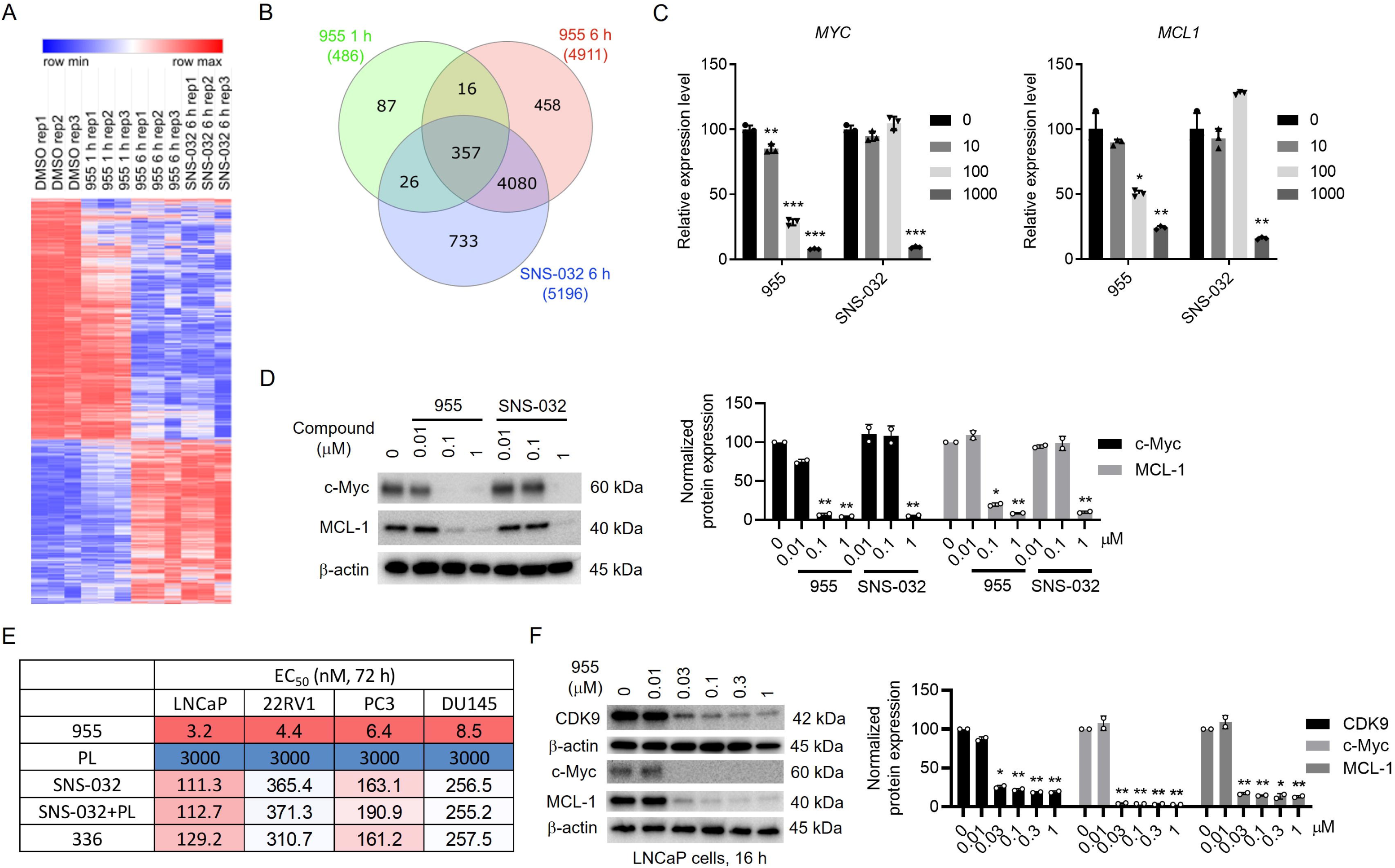
955 is a more potent antitumor agent than SNS-032. (A) Heatmap and (B) Venn diagram to show the differentially expressed genes in MOLT4 cells after **955** and SNS-032 treatments; (C) and (D) to demonstrate that the effects of **955** and SNS-032 on the expression of the two important CDK9 downstream targets MYC and MCL1 in MOLT4 cells at the levels of transcription and translation determined by qPCR and immunoblot, respectively. The qPCR data are expressed as mean ± SD of three biological replicates. Statistical significance was calculated with unpaired two-tailed Student’s *t*-tests. \**P* < 0.05; \*\**P* < 0.01; \*\*\**P* < 0.001. (E) **955** exhibits higher potencies against multiple prostate cancer cells compared with PL, SNS-032, SNS-032 plus PL, and **336**. The data are presented as a mean from three independent experiments; (F) **955** induces CDK9 degradation and downregulates the expression of c-Myc and MCL-1 in LNCaP prostate cancer cells. In panels D and F, representative immunoblots are shown and β-actin was used as an equal loading control. The quantification of the relative CDK9, c-Myc, or MCL-1 protein content in the immunoblots is presented as mean ± SD (*n* = 2 biologically independent experiments) in the bar graph (right panel). Statistical significance was calculated with unpaired two-tailed Student’s *t*-tests. \**P* < 0.05; \*\**P* < 0.01. See also Figures S5, S6 and Table S4.

### The PL-Ceritinib conjugate degrades ALK-fusion oncoprotein

To further evaluate the potential of PL as a novel covalent KEAP1 ligand to generate PROTACs for the degradation of oncoproteins, we synthesized several PL-ceritinib conjugates to test if these compounds can degrade EML4-ALK, an oncogenic fusion protein in NSCLC (Figure 6A) because EML4-ALK-positive NSCLC patients frequently develop resistance to the anaplastic lymphoma kinase (ALK) inhibitor ceritinib.^45^ In addition, ceritinib has been used for generating a CRBN-ALK PROTAC.^46^ Here we showed one of those conjugated compounds, **819**, can degrade EML4-ALK in a concentration-dependent manner in NCI-H2228 NSCLC cells (Figure 6B). The degradation of EML4-ALK can be blocked by the pre-treatment of the cells with MG132, MLN4924, and DMF (Figure 6C, D), suggesting that **819** may also recruit KEAP1 to mediate the degradation of EML4-ALK in NCI-H2228 cells in a UPS-dependent manner. With further optimization of the linker, we expect that PL-ceritinib conjugates can more potently degrade EML4-ALK in NSCLC cells.

**Figure 6.**
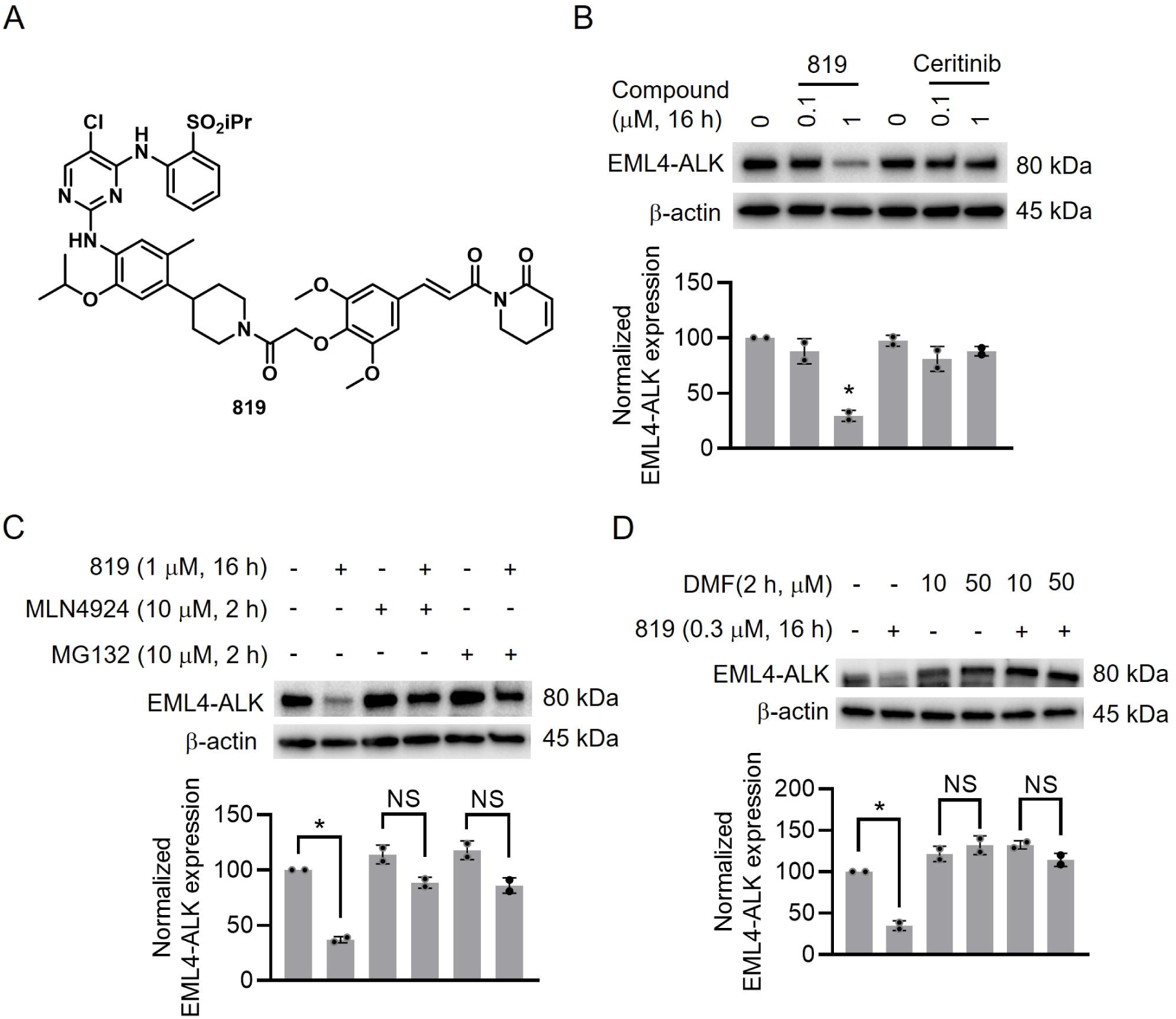
The PL-Ceritinib conjugate degrades the EML4-ALK-fusion protein. (A) The structure of PL-ceritinib conjugate **819**; (B) **819** but not ceritinib degrades EML4-ALK in NCI-H2228 cells; (C) Pre-treatment with the neddylation inhibitor MLN4924 or proteasome inhibitor MG132 blocks EML4-ALK degradation induced by **819;** and (D) Pre-treatment with the KEAP1 inhibitor DMF blocks the **819**-induced EML4-ALK degradation. In panels B, C and D, representative immunoblots are shown and β-actin was used as an equal loading control. The quantification of the relative EML4-ALK protein content in the immunoblots is presented as mean ± SD (*n* = 2 biologically independent experiments) in the bar graph (bottom panel). Statistical significance was calculated with unpaired two-tailed Student’s *t*-tests. \**P* < 0.05; NS: not significant.

### Limitations of the study

Our present data suggest that **955** should covalently bind KEAP1 because compound **336** lacking the two Michael acceptors (acrylamide groups) does not induce CDK9 degradation (Figure 2G, H) and **336** also loses the ability to form a strong ternary complex with CDK9 and KEAP1 as shown by nanoBRET assay (Figure 3I). Previous studies have shown that CDDO-Me binds to Cys151 of KEAP1. According to the results in Figure 3D that CDDO-Me/IM can compete with **955** to bind to KEAP1, it is highly possible that **955** also binds to Cys151 on the BTB domain of KEAP1. However, we cannot exclude the possibility that **955** may have the ability to recruit KEAP1 by binding to other cysteines. Additionally, further studies are needed to determine which Michael acceptor plays a major role in binding to KEAP1. TurboID-bait assay is very useful for identifying E3 ligases for novel protein degraders. In the present work, we use a transient overexpression system to perform this assay. It will be ideal to perform this assay in cells stably expressing TurboID-CDK9 with endogenous CDK9 deletion.

### Significance

Despite the human proteome encoding more than 600 E3 ligases, only a few E3 ligases have been used for PROTAC design due to the limited availability of E3 ligase ligands. Therefore, finding new E3 ligase ligands to expand the toolbox of PROTAC technology is critical for the further development of this field. Here, we design and synthesize a series of PL-SNS-032 conjugates and find that the lead compound of PL-SNS-032 conjugates, **955**, potently induces CDK9 degradation in a UPS-dependent manner. To find the E3 ligase(s) recruited by **955**, we use the TurboID-bait assay to identify the proteins that transiently interact with CDK9 induced by **955**. We found that KEAP1 is the CDK9-degrading E3 ligase recruited by **955** through covalent binding of PL, which is further confirmed genetically by siRNA and CRISPR-Cas9 knock down/out and rescue assays. Furthermore, EML4-ALK protein can also be successfully degraded by the PL-ceritinib conjugate **819**, which provides additional evidence to demonstrate that PL can be used as a KEAP1 recruiter to induce the degradation of different POIs. Compared with the other two commonly used E3 ligases (VHL and CRBN), KEAP1 is highly expressed in many tumor cells such as lung, kidney, breast, prostate, and brain cancer cells. Therefore, PL-based PROTACs may achieve better degradation efficacy in those cancer cells. Additionally, with a relatively smaller molecular size (MW=317.3 Da) of PL, PL-based PROTACs might possess more favorable physicochemical properties for drug development after further characterization of the binding mode of PL with KEAP1 and optimization to improve its specificity. Taken together, our study demonstrates that natural products are an important source for the discovery of new E3 ligase ligands and TurboID-bait assay is a powerful tool to identify E3 ligases recruited by natural compounds and novel E3 ligase ligands.

## ACKNOWLEDGMENTS

This study was supported in part by US National Institutes of Health (NIH) grants R01 CA242003 (D.Z. and G.Z.), R01 CA241191 (G.Z., M.K., and D.Z.), R01 AG063801 (D.Z. and G.Z.) and R56AG065635 (G.Z.). A Mays Cancer Center Early Career Pilot Award from CCSG (P30 CA054174) (D.L.). A Leukemia and Lymphoma Society Specialized Center of Research (J.D.L.) and LLS Special Fellow Award (A.S.). We thank the Mass Spectrometry Research and Education Center and the funding source: NIH S10 OD021758-01A1. The authors would like to thank Ms. Janet Wiegand and Alexandra M. Fahnlander for editing the manuscript.

## AUTHOR CONTRIBUTIONS

Conceptualization, G.Z., D.Z., and D.L.; chemistry, Y.X., X.L., W.H., X.Z., and G.Z.; cell biology, J.P., A.S., Y.Y., N.H., Q.Y., J.D.L., and D.L.; biochemistry, J.P., S.Z., and D.L.; proteomics, J.P., S.G.M., K.B.B., M.K., and D.L.; manuscript draft, J.P., Y.X., X.L., G.Z., D.Z., and D.L. All authors contributed to the review of the manuscript.

## Main tables and legends

N/A

## STAR METHODS

Detailed methods are provided in the online version of this paper and include the following:

- KEY RESOURCES TABLE
- LEAD CONTACT
- MATERIALS AVAILABILITY
- DATA AND CODE AVAILABILITY
- EXPERIMENTAL MODEL AND SUBJECT DETAILS

- Cell Lines
- METHOD DETAILS

- Immunoblotting
- Viability assay
- Competitive ABPP and LC-MS/MS
- TurboID-bait and LC-MS/MS
- TMT-based proteomics
- RNA-seq
- Reverse transcription and quantitative PCR (RT-qPCR)
- CRISPR knockout of KEAP1
- KEAP1 rescue assay
- Expression and purification of recombinant KEAP1
- Gel-based ABPP assay
- NanoBRET ternary complex formation assay
- Chemistry Methods
- QUANTIFICATION AND STATISTICAL ANALYSIS

## Supplementary Tables

**Table S1. The PL-binding proteins identified by competitive ABPP combined with LC-MS/MS (related to Figure S1 and STAR Methods).**

See accompanying Excel file.

**Table S2. The 955 binding proteins identified by TurboID-bait combined with LC-MS/MS (related to Figure 3 and STAR Methods)**.

See accompanying Excel file.

**Table S3. The TMT proteomics results in MOLT4 cells under 955 or SNS-032 treatment (related to Figure 4 and Figure S5).**

See accompanying Excel file.

**Table S4. The RNA-seq results in MOLT4 cells under 955 or SNS-032 treatment (related to Figure 5 and Figure S5).**

See accompanying Excel file.

## LEAD CONTACT

Requests for resources and reagents should be directed to the Lead Contact: Dongwen Lv.

## MATERIALS AVAILABILITY

The reagents generated in this study will be made available on request with a completed Materials Transfer Agreement.

## DATA AND CODE AVAILABILITY

- The authors declare that all data supporting the findings of this study are available within the paper and its supporting information files. RNA sequencing data supporting the current study have been deposited to Gene Expression Omnibus with accession number GEO: GSE206612. The proteomics data generated in this study are available at PRIDE archive with accession numbers PXD039385 and PXD039401.
- This paper does not report original code.
- Any additional information required to reanalyze the data reported in this paper is available from the lead contact upon request. Data used in this study are available via the Lead Contact upon reasonable request.

## EXPERIMENTAL MODEL AND SUBJECT DETAILS

### Cell Lines

Human T-ALL MOLT4 (Cat. No. CRL-1582) (donor sex: male), K562 (Cat. No. CCL-243) (donor sex: female), NCI-H1299 (Cat. No. CRL-5803) (donor sex: male), NCI-H2228 (Cat. No. CRL-5935) (donor sex: female), LNCaP (Cat. No. CRL-1740) (donor sex: male), 22RV1 (Cat. No. CRL-2505) (donor sex: male), PC3 (Cat. No. CRL-1435) (donor sex: male), DU145 (Cat. No. HTB-81) (donor sex: male), and epithelial kidney HEK 293T (293T, Cat. No. ACS-4500) (donor sex: female) cell lines were recently purchased from American Type Culture Collection (ATCC, Manassas, VA, USA). MOLT4, K562, NCI-H1299, NCI-H2228, LNCap, and 22RV1 cell lines were cultured in RPMI 1640 medium (Cat No. 22400-089, Thermo Fisher Scientific, Waltham, MA, USA) supplemented with 10% (v/v) heat-inactivated fetal bovine serum (FBS, Cat. No. S11150H, Atlanta Biologicals, Flowery Branch, GA, USA), 100 U/mL penicillin and 100 µg/mL streptomycin (Pen-Strep, Cat. No. 15140122, Thermo Fisher Scientific). PC3 cells were cultured in complete F-12K medium (Kaighn’s modification of Ham’s F-12 medium, Cat. No. 30-2004, ATCC) supplemented with 10% (v/v) heat-inactivated FBS, 100 U/mL penicillin, and 100 µg/mL streptomycin. DU145 cells were cultured in complete Eagle’s Minimum Essential Medium (EMEM, Cat. No. 30-2003, ATCC) supplemented with 10% (v/v) heat-inactivated FBS, 100 U/mL penicillin, and 100 µg/mL streptomycin. 293T cells were cultured in complete Dulbecco’s modified Eagle medium (DMEM, Cat. No. 12430054, Thermo Fisher Scientific) with 10% (v/v) heat-inactivated FBS, 100 U/mL penicillin, and 100 µg/mL streptomycin. All the cell lines were maintained in a humidified incubator at 37 °C and 5% CO_2._

NEB® 5-alpha Competent E. coli cells (Cat. No. C2987H, NEB) were cultured in LB broth at 37 °C to produce plasmids for the TurboID-bait assay. One Shot™ Stbl3™ Chemically Competent E. coli cells (Cat. No. C737303, Fisher Scientific) were cultured in LB broth at 37 °C to produce KEAP1 knockout plasmids. BL21(DE3) Chemically Competent E. coli cells (Cat. No. C600003, Fisher Scientific) were cultured in LB broth at 37 °C to express His-KEAP1 proteins.

## METHOD DETAILS

### Immunoblotting

Cells were collected and lysed in RIPA lysis buffer (Cat. # BP-115DG, Boston Bio Products, Ashland, MA, USA) supplemented with protease and phosphatase inhibitor cocktails (Cat. # PPC1010, Sigma-Aldrich, St. Louis, MO, USA). Protein samples were made and immunoblotting was performed as described previously^4,6^. Antibodies purchased from Cell Signaling Technologies (CST) and the dilutions are as follows: Biotin (Cat. No. 5571S, 1:1000), PARP (Cat. No. 9542S, 1:1000), c-Myc (Cat. No. 5605S, 1:1000), MCL1 (Cat. No. 5453S, 1:1000), and ALK (Cat. No. 3791S, 1:1000), TRAF6 (Cat. No. 8082S, 1:1000). KEAP1 (Cat. No. 10503-2-AP, 1:1000) and NRF2 (Cat. No. 16396-1-AP, 1:1000) antibodies were purchased from Proteintech (Cat. No. 10503-2-AP, 1:1000). CDK9 antibody was purchased from Santa Cruz (Cat. No. sc-13130, 1:500). V5-tag antibody was purchased from Bethyl (Cat. No. A190-120P, 1:1000). TRIP12 antibody was purchased from Fortis (Cat. No. A301-814A-M, 1:1000). β-actin antibody was purchased from MP Biomedicals (Cat. No. 8691001, 1:20 000).

### Viability assay

Viability (MTS) assays in various cancer cells followed the methods described in our previous report.^4^ In general, cancer cells in cell culture medium were seeded in 96-well plates (100 µl per well) at the optimized densities (50,000–100,000 suspension cells, 3,000–5,000 adherent cells). Suspension cells were treated 30 min after seeding, whereas adherent cells were allowed to adhere overnight and then treated with compounds. Compounds were prepared in cell culture medium and 100 µl of 2× treatment-containing medium was added to each well. Cell culture medium without treatment was added in control wells, and wells containing medium without cells served as background. The outer wells of the 96-well plate were not used for testing but filled with 250 µl of PBS to reduce the evaporation of medium from the inner wells. Each compound was tested at nine different concentrations with three replicates. The cell viability was measured after 72 h by tetrazolium-based MTS assay. MTS reagent (2 mg/ml stock; Cat. No. G1111, Promega) was freshly supplemented with phenazine methosulfate (PMS, 0.92 mg/ml stock, Cat. No. P9625, Sigma-Aldrich) in 20:1 ratio, and 20 µl of this mixture was added to each control and treated well. The cells were incubated at 37 °C and 5% CO_2_ for 4 h, and then the absorbance was recorded at 490 nm using Biotek’s Synergy Neo2 multimode plate reader (Biotek). The average absorbance value of background wells was subtracted from the absorbance value of each control and treatment well, and percent cell viability ((At / A0) × 100)) was determined in each treatment well, where At is the absorbance value of the treatment well and A0 is the average absorbance value of control wells after background subtraction. The data were expressed as average percentage cell viability and fitted in non-linear regression curves using GraphPad Prism 9 (GraphPad Software, La Jolla, CA, USA). For measuring the cell-killing effects in KEAP1 siRNA- and control siRNA-treated cells, LNCaP and H1299 cells were treated with control siRNA or siRNAs targeting KEAP1 for 48 h. Then cells were reseeded in 96-well plate and allowed to adhere overnight, then treated with indicated compound for 24 h. The viability was measured as described above.

### Competitive ABPP and LC-MS/MS

#### Competitive ABPP

MOLT4 cells (2 ×10^7^) were pre-treated with DMSO or 20 μM PL for 4 h and then treated with 1 μM PL-Alkyne probe^32^ for an additional 4 h. The cells were then harvested and lysed in 1x Dulbecco’s phosphate-buffered saline (PBS) buffer (Cat. # D1408, Sigma-Aldrich) containing protease and phosphatase inhibitors. CuAAC reaction was performed as previously described^23^ and biotinylated proteins were enriched using Pierce™ streptavidin agarose beads (Cat. # 20353, ThermoFisher).

#### Sample preparation and LC-MS/MS analysis

Protein samples were reduced, alkylated, and digested on-bead using filter-aided sample preparation^47^ with sequencing grade modified porcine trypsin (Cat. No. V5111, Promega). Nano-liquid chromatography tandem mass spectrometry (Nano-LC/MS/MS) was performed on a Thermo Scientific Q Exactive HF Orbitrap mass spectrometer equipped with an EASY Spray nanospray source (Thermo Scientific) operated in positive ion mode. The LC system was an UltiMate™ 3000 RSLCnano system (Thermo Scientific). The mobile phase A was water containing 0.1% formic acid and the mobile phase B was acetonitrile with 0.1 % formic acid. The mobile phase A for the loading pump was water containing 0.1 % trifluoracetic acid. 5 μL of sample is injected onto a PharmaFluidics μPAC^TM^ C18 trapping column (C18, 5 μm pillar diameter, 10 mm length, 2.5 μm inter-pillar distance). at 10 μL/ml flow rate. This was held for 3 min and washed with 1 % B to desalt and concentrate the peptides. The injector port was switched to inject, and the peptides were eluted off of the trap onto the column. PharmaFluidics 50 cm μPAC^TM^ was used for chromatographic separations (C18, 5 μm pillar diameter, 50 cm length, 2.5 μm inter-pillar distance). The column temperature was maintained 40 °C. A flowrate of 750 nl/min was used for the first 15 min and then the flow was reduced to 300 nl/min. Peptides were eluted directly off the column into the Q Exactive system using a gradient of 1% B to 20% B over 100 minutes and then to 45% B in 20 minutes for a total run time of 150 minutes:

| **Time (min)** | **% B** | **Flow Rate (nL/min)** |
| --- | --- | --- |
| 0 | 1 | 750 |
| 3 | 1 | 750 |
| 15 | 5 | 750 |
| 15.1 | 5 | 300 |
| 100 | 20 | 300 |
| 123 | 45 | 300 |
| 130 | 95 | 300 |
| 135 | 95 | 300 |
| 135.1 | 1 | 300 |
| 150 | 1 | 300 |

The MS/MS was acquired according to standard conditions established in the lab. The EASY Spray source operated with a spray voltage of 1.5 KV and a capillary temperature of 200 °C. The scan sequence of the mass spectrometer was based on the original TopTen™ method; the analysis was programmed for a full scan recorded between 375– 1575 Da at 60,000 resolution, and a MS/MS scan at resolution 15,000 to generate product ion spectra to determine amino acid sequence in consecutive instrument scans of the fifteen most abundant peaks in the spectrum. The AGC Target ion number was set at 3e6 ions for full scan and 2e5 ions for MS2 mode. Maximum ion injection time was set at 50 ms for full scan and 55 ms for MS2 mode. Micro scan number was set at 1 for both full scan and MS2 scan. The HCD fragmentation energy (N)CE/stepped NCE was set to 28 and an isolation window of 4 *m/z*. Singly charged ions were excluded from MS2. Dynamic exclusion was enabled with a repeat count of 1 within 15 seconds and to exclude isotopes. A Siloxane background peak at 445.12003 was used as the internal lock mass. HeLa protein digest standard is used to evaluate the integrity and the performance of the columns and mass spectrometer. If the number of protein IDs from the HeLa standard falls below 2700, the instrument is cleaned, and new columns are installed.

#### Data processing and analysis

All MS/MS samples were analyzed using Sequest (Thermo Fisher Scientific, San Jose, CA, USA; version IseNode in Proteome Discoverer 2.4.0.305). Sequest was set up to search Homo sapiens (NcbiAV TaxID=9606) (v2017-10-30) assuming the digestion enzyme trypsin. Sequest was searched with a fragment ion mass tolerance of 0.020 Da and a parent ion tolerance of 10.0 ppm. Carbamidomethyl of cysteine was specified in Sequest as a fixed modification. Met-loss of methionine, met-loss+Acetyl of methionine, oxidation of methionine and acetyl of the N-terminus were specified in Sequest as variable modifications. Protein identifications were accepted if they could be established with less than 1.0% false discovery and contained at least 2 identified peptides.

### TurboID-bait and LC-MS/MS

#### Construction of the pcDNA3-V5-TurboID-CDK9 plasmid

To build the TurboID-bait assay, we first constructed a V5-TurboID-tagged CDK9 plasmid (pDL2518) through Gibson assembly method. The template plasmids HA-CDK9 (a gift from Andrew Rice, plasmid # 28102) and V5-TurboID-NES_pCDNA3 (a gift from Alice Ting, plasmid #107169) were purchased from Addgene. The primer pair (5′- cagtctgcggtctgccgaaaagctgcagATGGCAAAGCAGTACGACTCGGTGGAGT-3′ and 5′-cagggtcaggcgctccaggggaggcagTCAGAAGACGCGCTCAAACTCCGTCTGGT-3′) and the template HA-CDK9 were used to clone the CDK9 fragment. The two primer pairs: pair-1 (5′- ACCAGACGGAGTTTGAGCGCGTCTTCTGActgcctcccctggagcgcctgaccctg-3′ and 5′-caatgatgagcacttttaaagttctgctatgtggcgcggtattatcccgtattgac-3′); pair-2 (5′- gtcaatacgggataataccgcgccacatagcagaactttaaaagtgctcatcattg-3′ and 5′- ACTCCACCGAGTCGTACTGCTTTGCCATctgcagcttttcggcagaccgcagactg-3′) and the template V5-TurboID-NES_pCDNA3 were used to clone the V5-TurboID fragment. The PCR fragments were assembled using NEBuilder® HiFi DNA Assembly Master Mix (Cat. No. E2621, NEB, Ipswich, MA, USA). DNA sequences in all these plasmids were authenticated by automatic sequencing.

#### TurboID-bait assay

293T (6 x 10^5^ for WB detection and 1.2 x 10^6^ for MS) cells were transfected with plasmid V5-TurboID-CDK9 using Lipofectamine 2000 as previously described.^6^ After 36 h, the transfected cells were pretreated with 10 μM MG132 for 1 h and followed up with co-treatment of DMSO or compound **955** and 50 μM D-Biotin at 37 °C for 6 h. Then the cells were harvested and lysed in 1x PBS buffer (Cat. # D1408, Sigma-Aldrich). After centrifugation at 20,000 x g for 60 min at 4 °C, the supernatant soluble fraction was used for further analysis. The same amount of protein was used for the enrichment of biotinylated proteins using100 μl Pierce™ streptavidin agarose beads (Cat. # 20353, ThermoFisher) overnight at 4 °C. The beads were further washed 3 times with PBS and deionized water respectively for LC-MS/MS analysis or boiled at 95 °C for 5 min in 100 ul 1x Laemmli SDS-Sample buffer (Cat #. BP-110R, Boston BioProducts) for immunoblot analysis.

#### Sample preparation and LC-MS/MS analysis

Protein samples were reduced, alkylated, and digested on-bead as mentioned above. Tryptic peptides were then separated by reverse phase XSelect CSH C18 2.5 um resin (Waters) on an in-line 150 x 0.075 mm column using an UltiMate 3000 RSLCnano system (Thermo). Peptides were eluted using a 60 min gradient from 98:2 to 65:35 buffer A:B ratio (Buffer A: 0.1% formic acid, 0.5% acetonitrile; Buffer B: 0.1% formic acid, 99.9% acetonitrile). Eluted peptides were ionized by electrospray (2.4 kV) followed by mass spectrometric analysis on an Orbitrap Eclipse Tribrid mass spectrometer (Thermo). MS data were acquired using the FTMS analyzer in profile mode at a resolution of 120,000 over a range of 375 to 1200 *m/z*. Following HCD activation, MS/MS data were acquired using the ion trap analyzer in centroid mode and normal mass range with a normalized collision energy of 30%. Proteins were identified by database search using MaxQuant (Max Planck Institute) with a parent ion tolerance of 2.5 ppm and a fragment ion tolerance of 0.5 Da. Scaffold Q+S (Proteome Software) was used to verify MS/MS based peptide and protein identifications. Protein identifications were accepted if they could be established with less than 1.0% false discovery and contained at least 2 identified peptides. Protein probabilities were assigned by the Protein Prophet algorithm^48^ and to perform reporter ion-based statistical analysis.

### TMT-based proteomics

#### Sample Preparation and LC-MS/MS analysis

Total protein from cell pellets was reduced, alkylated, and purified by chloroform/methanol extraction prior to digestion with sequencing grade modified porcine trypsin (Cat. No. V5111, Promega). Tryptic peptides were labeled using tandem mass tag isobaric labeling reagents (Cat. No. A34808, ThermoFisher) following the manufacturer’s instructions and combined into one 11-plex sample group. The labeled peptide multiplex was separated into 46 fractions on a 100 x 1.0 mm Acquity BEH C18 column (Waters) using an UltiMate 3000 UHPLC system (Thermo) with a 50 min gradient from 99:1 to 60:40 buffer A:B (Buffer A: 0.1% formic acid, 0.5% acetonitrile; Buffer B: 0.1% formic acid, 99.9% acetonitrile. Both buffers adjusted to pH 10 with ammonium hydroxide for offline separation) ratio under basic pH conditions, and then consolidated into 18 super-fractions. Each super-fraction was then further separated by reverse phase XSelect CSH C18 2.5 um resin (Waters) on an in-line 150 x 0.075 mm column using an UltiMate 3000 RSLCnano system (Thermo). Peptides were eluted using a 75 min gradient from 98:2 to 60:40 buffer A:B ratio. Eluted peptides were ionized by electrospray (2.4 kV) followed by mass spectrometric analysis on an Orbitrap Eclipse Tribrid mass spectrometer (Thermo) using multi-notch MS3 parameters. MS data were acquired using the FTMS analyzer in top-speed profile mode at a resolution of 120,000 over a range of 375 to 1500 *m/z*. Following CID activation with normalized collision energy of 35.0, MS/MS data were acquired using the ion trap analyzer in centroid mode and normal mass range. Using synchronous precursor selection, up to 10 MS/MS precursors were selected for HCD activation with normalized collision energy of 65.0, followed by acquisition of MS3 reporter ion data using the FTMS analyzer in profile mode at a resolution of 50,000 over a range of 100-500 *m/z*.

#### Data processing and analysis

Proteins were identified and MS3 reporter ions quantified using MaxQuant (version 2.0.3.0; Max Planck Institute) against the Homo sapiens UniprotKB database (March 2021) with a parent ion tolerance of 3 ppm, a fragment ion tolerance of 0.5 Da, and a reporter ion tolerance of 0.003 Da. Scaffold Q+S (Proteome Software) was used to verify MS/MS based peptide and protein identifications (protein identifications were accepted if they could be established with less than 1.0% false discovery and contained at least 2 identified peptides; protein probabilities were assigned by the Protein Prophet algorithm^48^ and to perform reporter ion-based statistical analysis. Protein TMT MS3 reporter ion intensity values were assessed for quality using ProteiNorm for a systematic evaluation of normalization methods ^49^. Cyclic loess normalization^50^ was utilized since it had the highest intragroup correlation and the lowest variance amongst the samples. Statistical analysis was performed using Linear Models for Microarray Data (limma) with empirical Bayes (eBayes) smoothing to the standard errors.^50^ Proteins with an FDR adjusted p-value < 0.05 and a fold change > 2 were considered to be significant.

### RNA-seq

Total RNA was extracted using RNeasy Plus Mini Kit (Qiagen, Cat. No. 74134). The library construction, cluster generation and DNBseq (BGI) sequencing were performed with BGI following the previously reported methods.^51^ Raw fastq data were analyzed by using Galaxy (https://usegalaxy.org/) as described previously.^52^ Human genome (hg38) was used as the reference genome. Differentially expressed gene was analyzed by using DESeq2 based on a false-discovery rate–adjusted q-value (q< 0.05). Genes with more than 2-fold change were selected for further analysis.

### Reverse transcription and quantitative PCR (RT-qPCR)

Total RNA was extracted as mentioned above. Reverse transcription and quantitative PCR (qPCR) was performed as described in our previous study.^6^ Primers used for measuring gene transcriptional level: *GAPDH* primers (forward 5’- GACCACTTTGTCAAGCTCATTTC-3’ and reverse 5’- CTCTCTTCCTCTTGTGCTCTTG-3’) were described previously^32^. *MCL1* primers are forward 5’-ATCTCTCGGTACCTTCGGGAGC-3’ and reverse 5’- GCTGAAAACATGGATCATCACTCG-3’; *MYC* primers are forward 5’- GGCTCCTGGCAAAAGGTCA-3’ and reverse 5’-CTGCGTAGTTGTGCTGATGT-3’.

### CRISPR knockout of KEAP1

To knockout KEAP1, two lentiviral CRISPR knockout plasmids targeting human *KEAP1* gene were purchased from Abm (Cat. No. 254911110595). Packaging 293T cells were transfected with *KEAP1* single guide RNA (sgRNA) and helper vectors (pMD2.G and psPAX2; Addgene plasmid # 12259 and 12260) using Lipofectamine 3000 reagent (Cat. No. L3000015, Thermal Fisher). Medium containing lentiviral particles and 8 μg/mL polybrene (Sigma-Aldrich) was used to infect H1299 cells. Infected cells were selected in medium containing 2 μg/mL puromycin. The sgRNA target sequences are as follows: sg1: GTACGCCTCCACTGAGTGCA; sg2: TGACAGCACCGTTCATGACG. Single clones were selected by serial dilution.

### KEAP1 rescue assay

KEAP1-sg1 knockout H1299 (4 x 10^5^) cells were transfected with or without plasmid expressing Halo-KEAP1 (Addgene, a gift from Yimon Aye, plasmid # 58240) using Lipofectamine 3000 reagent (Cat. No. L3000015, Thermal Fisher). After 24 h, the cells in each well of 6-well plate were divided into two wells and cultured for additional 16 h. The cells were then treated with DMSO or 0.1 μM **955** at 37 °C for 6 h, harvested and lysed for WB analysis.

### Expression and purification of recombinant KEAP1

The plasmid pet28a-His6-Keap1 for expressing His-KEAP1 was purchased from Addgene (a gift from Yimon Aye, plasmid # 62454). To purify His-KEAP1, the plasmid transformed BL21(DE3) Chemically Competent E. coli cells (Cat. No. C600003, Fisher Scientific) were grown in LB broth at 37 °C with shaking until the optical density at 600 nm reached 0.6-0.7. Isopropyl-β-D-thiogalactopyranoside (1 mM) was added to induce protein expression overnight at 16 °C. Cell pellet was resuspended in lysis buffer [50 mM Tris-Hcl (pH 7.5), 150 mM NaCl, 1 mM EDTA, 0.05% NP-40, 1 mg/ul Lysozyme, 1x protease inhibitor, and 1 mM DTT] and lysed using sonication. The lysate was cleared by centrifugation at 20,000 x g for 30 min at 4 °C and purified by using GE Healthcare His GraviTrap™ Kit (Cat. No. 45-002-016, Fisher Scientific). After elution, the proteins were concentrated by using Pierce Protein Concentrators PES, 30K MWCO (Cat. No. 88502, ThermoFisher) and then dialyzed into the buffer containing 50 mM HEPES, 150 mM NaCl, and 0.1 mM EDTA (pH 7.5).

### Gel-based ABPP assay

A gel-based ABPP assay was performed as previously described with some modifications.^25^ Briefly, Purified His-KEAP1 (0.2 μg) was diluted into 50 μL of PBS and 0.5 μL of either DMSO (vehicle) or indicated compound was added to incubate at 25 °C for 1 h, then the samples were treated with 20 μM IA-Alkyne (Cat. No. 7015, Tocris, prepared in DMSO) at 25 °C for 1 h. CuAAC reaction was performed following the protocol from ClickChemistryTools (Cat. No. 1001) and Azide-fluor 488 (Cat. No. 760765, Sigma) was used to react with IA-Alkyne. Protein pellet was dissolved in RIPA buffer and then 20 μL of 4× reducing Laemmli SDS sample loading buffer (Cat. No. J63615, Alfa Aesar) was added and heated at 95 °C for 5 min. The samples were resolved using precast 4–20% Tris-glycine gels (Mini-PROTEAN TGX, Cat. No. 4561094, Bio-Rad). Fluorescent imaging was performed on the ChemiDoc MP Imaging System (Bio-Rad). Then the gel was stained using the silver stain kit following the instructions from Thermo Fisher (Cat. No. 24612) and imaged using ChemiDoc.

### NanoBRET ternary complex formation assay

CMV HiBit (Cat. No. CS1956B03) was purchased from Promega. Plasmid HaloTag-KEAP1 (a gift from Yimon Aye, plasmid # 58240) was purchased from Addgene. HiBit-CDK9 were constructed through the Gibson assembly method. The primer pair (5′- TGGCTCGAGCGGTGGGAATTCTGGTatggcaaagcagtacgactcggtggag-3′ and 5′- TCTTCCGCTAGCTCCACCGGATCCTCCTCAgaagacgcgctcaaactccgtctggt-3′) and the template HA-CDK9 (a gift from Andrew Rice, plasmid # 28102) was used to clone the CDK9 fragment. The primer pair (5′- accagacggagtttgagcgcgtcttcTGAGGAGGATCCGGTGGAGCTAGCGGAAGA-3′ and 5′-ctccaccgagtcgtactgcttcgccatACCAGAATTCCCACCGCTCGAGCCA-3′) and pBit3.1-N (Cat. No. N2361, Promega) were used to clone the vector-containing fragment. The PCR fragments were assembled using NEBuilder® HiFi DNA Assembly Master Mix. DNA sequences in all these plasmids were authenticated by automatic sequencing. 293T cells (8 ×10^5^) were transfected with Lipofectamine 2000 (Life Technologies) and 1 μg HaloTag-KEAP1, 10 ng HiBit-CDK9 and 10 ng LgBit. After 24 h, 2 × 10^4^ transfected cells were seeded into white 96-well tissue culture plates in Gibco™ Opti-MEM™ I Reduced Serum Medium, No Phenol Red (Cat. No. 11-058-021, Fisher) containing 4% FBS with or without HaloTag NanoBRET 618 Ligand (Cat. No. PRN1662, Promega) and incubated overnight at 37 °C, 5% CO_2_. The following day, serially diluted compounds were added into the medium and plates were incubated at 37 °C, 5% CO_2_, for 6 h. After treatment, NanoBRET Nano-Glo Substrate (Cat. No. N1662, Promega) was added into the medium, and the contents were mixed by shaking the plate for 30 s before measuring donor and acceptor signals on Biotek plate reader. Dual-filtered luminescence was collected with a 450/50 nm bandpass filter (donor, NanoBiT-CDK9 protein) and a 610-nm longpass filter (acceptor, HaloTag NanoBRET ligand) using an integration time of 0.5 s. mBRET ratios were calculated following the NanoBRET™ Nano-Glo® Detection System (Cat. No. N1662, Promega).

### Chemistry Methods

General Methods. DMF and DCM were obtained via a solvent purification system by filtering through two columns packed with activated alumina and 4 Å molecular sieve, respectively. Water was purified with a Milli-Q Simplicity 185 Water Purification System (Merck Millipore). All other chemicals and solvents obtained from commercial suppliers were used without further purification. PL-Alkyne was obtained from our previous study.^32^ Flash chromatography was performed using silica gel (230–400 mesh) as the stationary phase. Reaction progress was monitored by thin-layer chromatography (silica-coated glass plates) and visualized by 256 nm and 365 nm UV light, and/or by LC-MS. ^1^ H NMR spectra were recorded in CDCl_3_ or CD_3_OD at 600 MHz, and ^13^C NMR spectrum were recorded at 151 MHz. Chemical shifts δ are given in ppm using tetramethylsilane as an internal standard. Multiplicities of NMR signals are designated as singlet (s), doublet (d), doublet of doublets (dd), triplet (t), quartet (q), A triplet of doublets (td), A doublet of triplets (dt), multiplet (m), and broad (br). All final compounds for biological testing were of ≥95.0% purity as analyzed by LC-MS, performed on an Advion AVANT LC system with the expression CMS using a Thermo Accucore™ Vanquish™ C18+ UHPLC Column (1.5 µm, 50 x 2.1 mm) at 40 °C. Gradient elution was used for UHPLC with a mobile phase of acetonitrile and water containing 0.1% formic acid. High resolution mass spectra (HRMS) were recorded on an Agilent 6230 Time-of-Flight (TOF) mass spectrometer.

### Synthetic procedures and characterizations

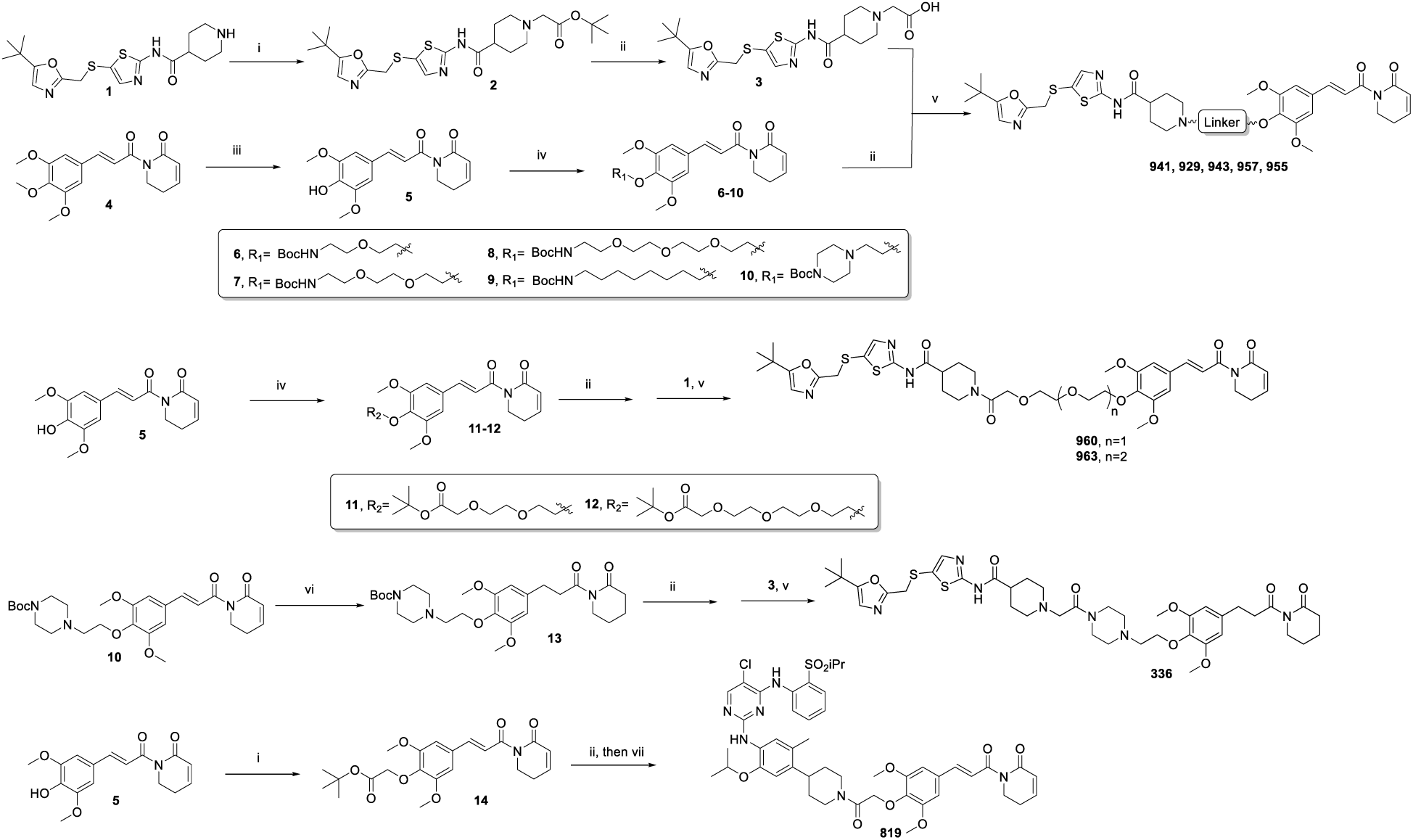

**Scheme S1.** Synthesis of PROTAC molecules. i) BrCH_2_COOtBu, K_2_CO_3_, DMF, r.t., overnight; ii) TFA, DCM, r.t., 5 h.; iii) AlCl_3_, THF, 0°C to r.t., 1 h; iv) 1,1′- (Azodicarbonyl)dipiperidine, tributylphosphine, alcohol, toluene, 0°C to r.t., overnight; v) HATU, DIPEA, DCM, r.t., overnight.; vi) H_2_, Pd/C, MeOH, overnight; vii) HATU, TEA, Ceritinib, DCM, r.t., overnight.

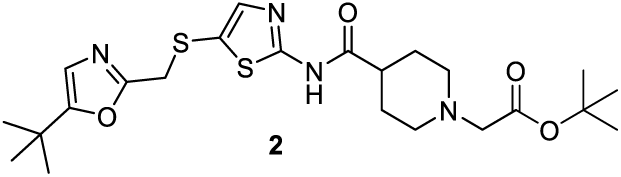

### tert-butyl 2-(4-((5-(((5-(tert-butyl)oxazol-2-yl)methyl)thio)thiazol-2- yl)carbamoyl)piperidin-1-yl)acetate (2)

To a solution of N-(5-(((5-(tert-butyl)oxazol-2-yl)methyl)thio)thiazol-2-yl)piperidine-4-carboxamide **1** (0.21 g, 0.55 mmol) in DMF (5 mL) was added K_2_CO_3_ (0.15 g, 1.1 mmol) and tert-Butyl bromoacetate (0.112 g, 0.58 mmol), the reaction mixture was stirred at room temperature overnight. Water was added and the mixture was extracted with EA (30 mL). The organic layer was washed with brine (30 mL*2), dried over anhydrous Na_2_SO_4_, filtered and concentrated under reduced pressure. The resulting crude was purified by flash chromatography (DCM/MeOH=50/1 to 25/1) to afford the compound **2** (0.586 g, 1.93 mmol, 61% yield) as a white solid. ^1^H NMR (600 MHz, Chloroform-*d*) δ 11.72 (s, 1H), 7.35 (s, 1H), 6.61 (s, 1H), 3.98 (s, 2H), 3.17 (s, 2H), 3.07 – 2.99 (m, 2H), 2.44 – 2.38 (m, 1H), 2.30 (td, *J* = 11.6, 2.6 Hz, 2H), 2.01 – 1.87 (m, 4H), 1.48 (s, 9H), 1.25 (s, 9H). ESI^+^, m/z 495.3 [M+H] ^+^.

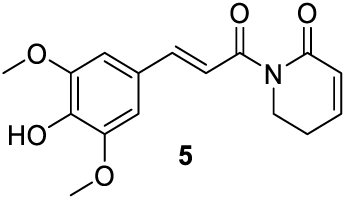

### (E)-1-(3-(4-hydroxy-3,5-dimethoxyphenyl)acryloyl)-5,6-dihydropyridin-2(1H)- one (5)

To a solution of (E)-1-(3-(3,4,5-trimethoxyphenyl)acryloyl)-5,6-dihydropyridin-2(1H)- one **4** (1.0 g, 3.15 mmol) in DCM (45 mL) was added AlCl_3_ (4.2 g, 31.5 mmol) portion-wise under ice bath, and the resulting reaction mixture was warmed to room temperature and stirred for 1 hr. Saturated NaHCO_3_ solution was added and the mixture was extracted with EA (60 mL*2). The combined organic layers were washed with brine (50 mL), dried over anhydrous Na_2_SO_4_, filtered and concentrated under reduced pressure to afford a gum. The gum was purified by flash chromatography (DCM/EA=10/1 to 10/2) to afford the compound **5** (0.586 g, 1.93 mmol, 61% yield) as a light yellow solid. ^1^H NMR (600 MHz, Chloroform-*d*) δ 7.69 (d, *J* = 15.5 Hz, 1H), 7.41 (d, *J* = 15.5 Hz, 1H), 6.94 (dt, *J* = 9.7, 4.2 Hz, 1H), 6.83 (s, 2H), 6.05 (dt, *J* = 9.7, 1.9 Hz, 1H), 5.75 (s, 1H), 4.04 (t, *J* = 6.5 Hz, 2H), 3.93 (s, 6H), 2.51 – 2.44 (m, 2H). ESI^+^, m/z 304.1 [M+H] ^+^.

### General procedure A for compounds 6-13

To a solution of (E)-1-(3-(4-hydroxy-3,5-dimethoxyphenyl)acryloyl)-5,6-dihydropyridin-2(1H)-one **5** (1 equiv.) and alcohol (1.1 equiv.) in toluene was added tributylphosphine (1.4 equiv.) and then 1,1′-(Azodicarbonyl)dipiperidine (1.3 equiv.) under ice bath, and the resulting reaction mixture was warmed to room temperature and stirred overnight with inserted nitrogen. Water was added and the mixture was extracted with EA (20 mL*2). The combined organic layers were washed with brine (20 mL), dried over anhydrous Na_2_SO_4_, filtered and concentrated under reduced pressure. The resulting crude was purified by flash chromatography to afford the desired compound.

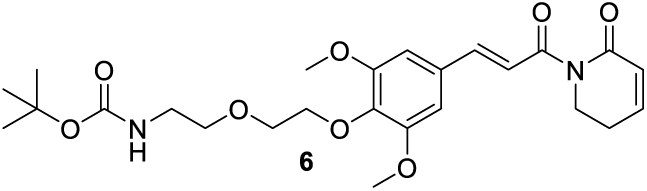

### tert-butyl (E)-(2-(2-(2,6-dimethoxy-4-(3-oxo-3-(6-oxo-3,6-dihydropyridin-1(2H)- yl)prop-1-en-1-yl)phenoxy)ethoxy)ethyl)carbamate (6)

84.2 mg (0.17 mmol, 75% yield) was obtained as colorless oil from **5** (70.0 mg, 0.23 mmol) and tert-butyl (2-(2-hydroxyethoxy)ethyl)carbamate (51.3 mg, 0.25 mmol) by using the general procedure A. ^1^H NMR (600 MHz, Chloroform-*d*) δ 7.67 (d, *J* = 15.5 Hz, 1H), 7.43 (d, *J* = 15.6 Hz, 1H), 7.00 – 6.90 (m, 1H), 6.81 (s, 2H), 6.04 (dt, *J* = 9.7, 1.8 Hz, 1H), 5.27 – 5.16 (m, 1H), 4.19 – 4.15 (m, 2H), 4.04 (t, *J* = 6.5 Hz, 2H), 3.89 (s, 6H), 3.77 – 3.73 (m, 2H), 3.60 (t, *J* = 5.0 Hz, 2H), 3.36 – 3.30 (m, 2H), 2.52 – 2.43 (m, 2H), 1.44 (s, 9H). ESI^+^, m/z 491.2 [M+H] ^+^

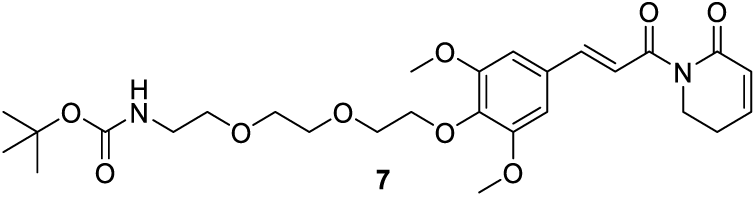

### tert-butyl (E)-(2-(2-(2-(2,6-dimethoxy-4-(3-oxo-3-(6-oxo-3,6-dihydropyridin- 1(2H)-yl)prop-1-en-1-yl)phenoxy)ethoxy)ethoxy)ethyl)carbamate (7)

89.5 mg (0.17 mmol, 64% yield) was obtained as colorless oil from **5** (80.0 mg, 0.26 mmol) and tert-butyl (2-(2-(2-hydroxyethoxy)ethoxy)ethyl)carbamate (72.3 mg, 0.29 mmol) by using the general procedure A. ^1^H NMR (600 MHz, Chloroform-*d*) δ 7.67 (d, *J* = 15.5 Hz, 1H), 7.42 (d, *J* = 15.5, 1.0 Hz, 1H), 6.98 – 6.93 (m, 1H), 6.80 (s, 2H), 6.04 (dq, *J* = 9.5, 1.5 Hz, 1H), 5.18 – 5.10 (m, 1H), 4.22 – 4.16 (m, 2H), 4.04 (t, *J* = 6.6 Hz, 2H), 3.87 (s, 6H), 3.82 – 3.78 (m, 2H), 3.73 – 3.69 (m, 2H), 3.65 – 3.62 (m, 2H), 3.58 – 3.53 (m, 2H), 3.36 – 3.27 (m, 2H), 2.51 – 2.46 (m, 2H), 1.42 (s, 9H). ESI^+^, m/z 535.4 [M+H]^+^

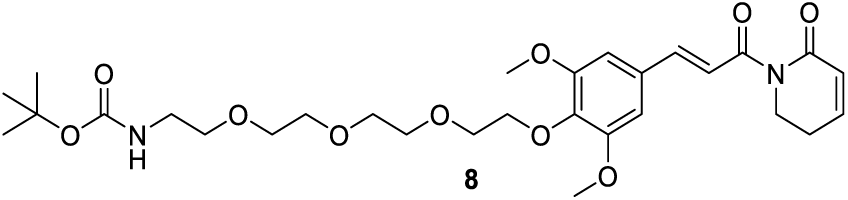

### tert-butyl (E)-(2-(2-(2-(2-(2,6-dimethoxy-4-(3-oxo-3-(6-oxo-3,6-dihydropyridin- 1(2H)-yl)prop-1-en-1-yl)phenoxy)ethoxy)ethoxy)ethoxy)ethyl)carbamate (8)

65.1 mg (0.11 mmol, 51% yield) was obtained as colorless oil from **5** (65.0 mg, 0.22 mmol) and tert-butyl (2-(2-(2-(2-hydroxyethoxy)ethoxy)ethoxy)ethyl)carbamate (70.4 mg, 0.24 mmol) by using the general procedure A. ^1^H NMR (600 MHz, Chloroform-*d*) δ 7.67 (d, *J* = 15.5 Hz, 1H), 7.42 (d, *J* = 15.5 Hz, 1H), 6.98 – 6.93 (m, 1H), 6.79 (s, 2H), 6.05 (dt, *J* = 9.7, 1.8 Hz, 1H), 5.17 – 5.04 (m, 1H), 4.20 – 4.16 (m, 2H), 4.04 (t, *J* = 6.5 Hz, 2H), 3.87 (s, 6H), 3.84 – 3.79 (m, 2H), 3.75 – 3.71 (m, 2H), 3.68 – 3.60 (m, 6H), 3.54 (t, *J* = 5.2 Hz, 2H), 3.34 – 3.28 (m, 2H), 2.48 (tdd, *J* = 6.3, 4.1, 1.9 Hz, 2H), 1.43 (s, 9H). ESI^+^, m/z 579.4 [M+H] ^+^

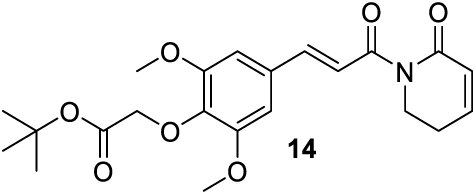

### tert-butyl (E)-2-(2,6-dimethoxy-4-(3-oxo-3-(6-oxo-3,6-dihydropyridin-1(2H)- yl)prop-1-en-1-yl)phenoxy)acetate (14)

40.2 mg (0.096 mmol, 85% yield) was obtained as colorless oil by using the same procedure as compound **2**. ^1^H NMR (600 MHz, Chloroform-*d*) δ 7.67 (d, *J* = 15.5 Hz, 1H), 7.42 (d, *J* = 15.5 Hz, 1H), 6.95 (d, *J* = 9.8 Hz, 1H), 6.79 (s, 2H), 6.05 (dd, *J* = 9.7, 1.9 Hz, 1H), 4.59 (s, 2H), 4.04 (t, *J* = 6.5 Hz, 2H), 3.87 (s, 6H), 2.50 – 2.45 (m, 2H), 1.47 (s, 9H). ESI^+^, m/z 418.3 [M+H] ^+^.

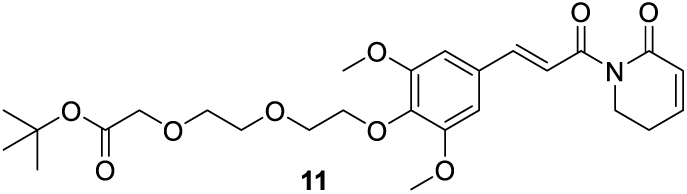

### tert-butyl (E)-2-(2-(2-(2,6-dimethoxy-4-(3-oxo-3-(6-oxo-3,6-dihydropyridin- 1(2H)-yl)prop-1-en-1-yl)phenoxy)ethoxy)ethoxy)acetate (11)

32.2 mg (0.064 mmol, 64% yield) was obtained as colorless oil from **5** (30 mg, 0.10 mmol) and tert-butyl 2- (2-(2-hydroxyethoxy)ethoxy)acetate (24.2 mg, 0.11 mmol) by using the general procedure A. ^1^H NMR (600 MHz, Chloroform-*d*) δ 7.67 (d, *J* = 15.6 Hz, 1H), 7.42 (d, *J* = 15.5 Hz, 1H), 6.98 – 6.93 (m, 1H), 6.79 (s, 2H), 6.05 (dt, *J* = 9.7, 1.8 Hz, 1H), 4.20 – 4.17 (m, 2H), 4.07 – 4.01 (m, 4H), 3.87 (s, 6H), 3.84 – 3.80 (m, 2H), 3.77 – 3.73 (m, 4H), 2.51 – 2.46 (m, 2H), 1.47 (s, 9H). ESI^+^, m/z 506.4 [M+H] ^+^.

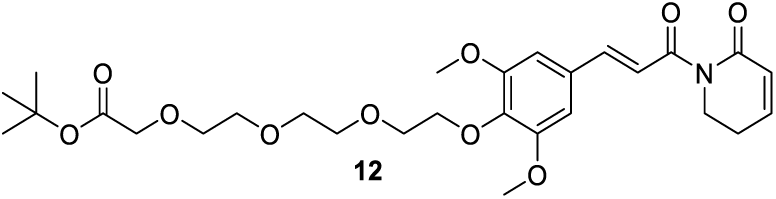

### tert-butyl (E)-2-(2-(2-(2-(2,6-dimethoxy-4-(3-oxo-3-(6-oxo-3,6-dihydropyridin- 1(2H)-yl)prop-1-en-1-yl)phenoxy)ethoxy)ethoxy)ethoxy)acetate (12)

71.5 mg (0.13 mmol, 48% yield) was obtained as colorless oil from **5** (83 mg, 0.27 mmol) and tert-butyl 2-(2-(2-(2-hydroxyethoxy)ethoxy)ethoxy)acetate (79.3 mg, 0.30 mmol) by using the general procedure A. ^1^H NMR (600 MHz, Chloroform-*d*) δ 7.67 (d, *J* = 15.5 Hz, 1H), 7.42 (d, *J* = 15.6 Hz, 1H), 6.99 – 6.92 (m, 1H), 6.79 (s, 2H), 6.05 (dt, *J* = 9.7, 1.8 Hz, 1H), 4.21 – 4.15 (m, 2H), 4.06 – 4.01 (m, 4H), 3.87 (s, 6H), 3.83 – 3.79 (m, 2H), 3.75 – 3.67 (m, 8H), 2.51 – 2.46 (m, 2H), 1.47 (s, 9H). ESI^+^, m/z 550.3 [M+H] ^+^.

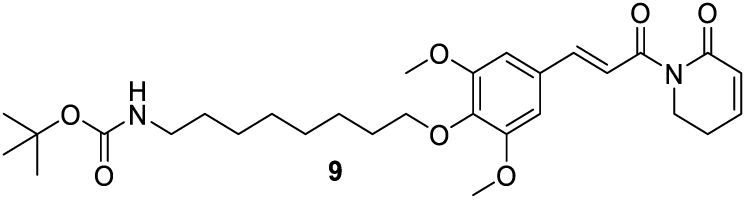

### tert-butyl (E)-(8-(2,6-dimethoxy-4-(3-oxo-3-(6-oxo-3,6-dihydropyridin-1(2H)- yl)prop-1-en-1-yl)phenoxy)octyl)carbamate (9)

31.9 mg (0.060 mmol, 60% yield) was obtained as colorless oil from **5** (30 mg, 0.10 mmol) and tert-butyl 2-(2-(2-(2- hydroxyethoxy)ethoxy)ethoxy)acetate (27.0 mg, 0.11 mmol) by using the general procedure A. ^1^H NMR (600 MHz, Chloroform-*d*) δ 7.68 (d, *J* = 15.5 Hz, 1H), 7.42 (d, *J* = 15.5 Hz, 1H), 6.98 – 6.92 (m, 1H), 6.80 (s, 2H), 6.05 (dt, *J* = 9.7, 1.8 Hz, 1H), 4.53 (s, 1H), 4.04 (t, *J* = 6.5 Hz, 2H), 3.99 (t, *J* = 6.8 Hz, 2H), 3.87 (s, 6H), 3.10 (q, *J* = 6.8 Hz, 2H), 2.52 – 2.46 (m, 2H), 1.75 – 1.71 (m, 2H), 1.50 – 1.26 (m, 19H). ESI^+^, m/z 531.3 [M+H] ^+^.

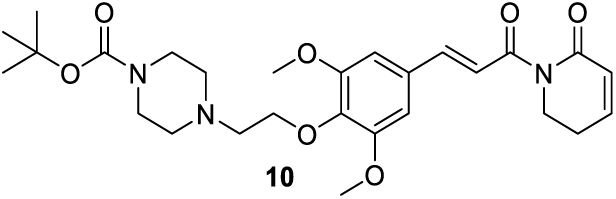

### tert-butyl (E)-4-(2-(2,6-dimethoxy-4-(3-oxo-3-(6-oxo-3,6-dihydropyridin-1(2H)- yl)prop-1-en-1-yl)phenoxy)ethyl)piperazine-1-carboxylate (10)

180.3 mg (0.35 mmol, 59% yield) was obtained as colorless oil from **5** (180.0 mg, 0.59 mmol) and tert-butyl 4-(2-hydroxyethyl)piperazine-1-carboxylate (149.5 mg, 0.65 mmol) by using the general procedure A. ^1^H NMR (600 MHz, Chloroform-*d*) δ 7.69 (d, *J* = 15.5 Hz, 1H), 7.44 (d, *J* = 15.5 Hz, 1H), 6.99 – 6.95 (m, 1H), 6.81 (s, 2H), 6.07 (dt, *J* = 9.7, 1.8 Hz, 1H), 4.14 (t, *J* = 5.7 Hz, 2H), 4.07 (t, *J* = 6.5 Hz, 2H), 3.88 (s, 6H), 3.48 (t, *J* = 5.0 Hz, 4H), 2.81 (t, *J* = 5.7 Hz, 2H), 2.59 – 2.48 (m, 6H), 1.48 (s, 9H). ESI^+^, m/z 516.2 [M+H]^+^.

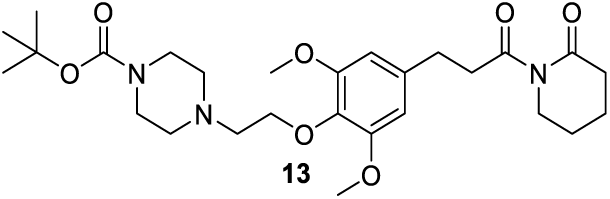

### tert-butyl 4-(2-(2,6-dimethoxy-4-(3-oxo-3-(2-oxopiperidin-1- yl)propyl)phenoxy)ethyl)piperazine-1-carboxylate (13)

To a solution of compound **10** (30.3 mg, 0.058 mmol) in 10 ml MeOH was added 10% Pd/C (6 mg, 20 wt % of **10**). After purged with nitrogen then hydrogen, the reaction was stirred at room temperature overnight. Filtered it and concentrated under reduced pressure. The resulting crude was purified by flash chromatography to afford the compound **13** (21.3 mg, 0.041 mmol, 71% yeild) as colorless oil. ^1^H NMR (600 MHz, Chloroform-*d*) δ 6.44 (s, 2H), 4.06 (t, *J* = 5.8 Hz, 2H), 3.81 (s, 6H), 3.71 (t, *J* = 5.9 Hz, 2H), 3.50 – 3.44 (m, 4H), 3.25 – 3.18 (m, 2H), 2.90 (t, *J* = 7.7 Hz, 2H), 2.78 (t, *J* = 5.8 Hz, 2H), 2.59 – 2.52 (m, 6H), 1.86 – 1.78 (m, 4H), 1.46 (s, 9H). ESI^+^, m/z 520.3 [M+H] ^+^.

### General procedure B for PROTACs

Boc-protected amine (1 equiv.) and tert-butyl protected acid (1 equiv.) were first converted to their corresponding free amine and acid by reacting with 2 mL trifluoroacetic acid in 2 mL DCM at room temperature for 4h. Each of them was then concentrated under vacuum to obtain the crude which was used without purification. To a solution of free amin and acid in 4 mL DCM was added DIPEA (10 equiv.) and HATU (1.1 equiv.). The reaction was stirred at room temperature overnight. Water was added and the mixture was extracted with EA (20 mL*2). The combined organic layers were washed with brine (15 mL), dried over anhydrous Na_2_SO_4_, filtered and concentrated under reduced pressure. The resulting mixture was purified by flash chromatography (DCM/MeOH=30/1 to 20/1) to afford the desired PROTACs.

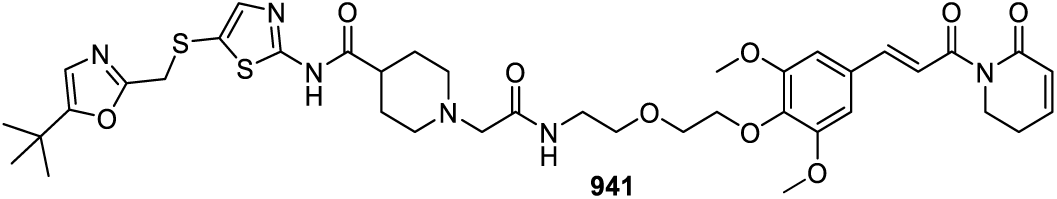

### (E)-N-(5-(((5-(tert-butyl)oxazol-2-yl)methyl)thio)thiazol-2-yl)-1-(2-((2-(2-(2,6- dimethoxy-4-(3-oxo-3-(6-oxo-3,6-dihydropyridin-1(2H)-yl)prop-1-en-1- yl)phenoxy)ethoxy)ethyl)amino)-2-oxoethyl)piperidine-4-carboxamide (941)

14.1 mg (0.017 mmol, 40% yield) was obtained as colorless gum from **6** (21.0 mg, 0.043 mmol) and **2** (21.3 mg, 0.043 mmol) by using the general procedure B. ^1^H NMR (600 MHz, Chloroform-*d*) δ 10.65 (br, 1H), 7.62 (d, *J* = 15.5 Hz, 1H), 7.47 (t, *J* = 6.1 Hz, 1H), 7.37 (d, *J* = 15.5 Hz, 1H), 7.30 (s, 1H), 6.99 – 6.92 (m, 1H), 6.77 (s, 2H), 6.59 (s, 1H), 6.07 (dt, *J* = 9.7, 1.8 Hz, 1H), 4.18 – 4.14 (m, 2H), 4.04 (t, *J* = 6.5 Hz, 2H), 3.96 (s, 2H), 3.86 (s, 6H), 3.80 – 3.74 (m, 2H), 3.64 (t, *J* = 5.1 Hz, 2H), 3.54 – 3.48 (m, 2H), 3.04 (s, 2H), 2.96 – 2.88 (m, 2H), 2.51 – 2.47 (m, 2H), 2.42 – 2.34 (m, 1H), 2.28 – 2.18 (m, 2H), 1.89 – 1.83 (m, 4H), 1.25 (s, 9H). ^13^C NMR (151 MHz, CDCl_3_) δ 172.59, 170.23, 168.87, 166.03, 161.88, 161.78, 158.84, 153.34, 145.65, 144.29, 143.54, 138.81, 130.79, 125.79, 121.29, 121.02, 120.11, 105.47, 77.24, 77.03, 76.82, 72.39, 70.22, 69.86, 61.67, 56.19, 53.13, 41.84, 41.70, 38.92, 34.96, 31.59, 28.57, 28.32, 24.81. MS (ESI); m/z: [M+H]^+^ calcd for C_39_H_51_N_6_O_9_S_2_^+^: 811.3153, found 811.3105.

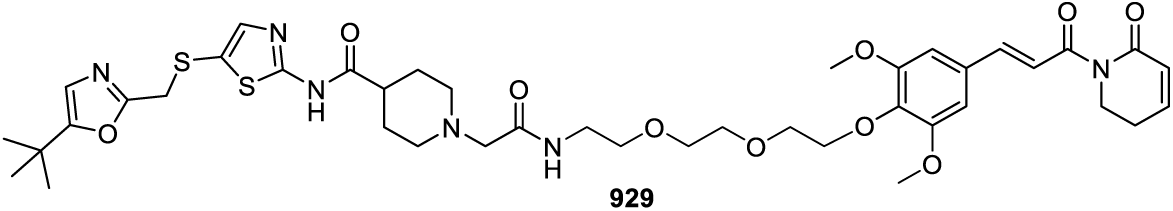

### (E)-N-(5-(((5-(tert-butyl)oxazol-2-yl)methyl)thio)thiazol-2-yl)-1-(2-((2-(2-(2-(2,6- dimethoxy-4-(3-oxo-3-(6-oxo-3,6-dihydropyridin-1(2H)-yl)prop-1-en-1- yl)phenoxy)ethoxy)ethoxy)ethyl)amino)-2-oxoethyl)piperidine-4-carboxamide (929)

12.1 mg (0.014 mmol, 42% yield) was obtained as colorless gum from **7** (18 mg, 0.034 mmol) and **2** (16.8 mg, 0.034 mmol) by using the general procedure B. ^1^H NMR (600 MHz, Chloroform-*d*) δ 10.47 (br, 1H), 7.64 (d, *J* = 15.5 Hz, 1H), 7.45 (t, *J* = 5.7 Hz, 1H), 7.39 (d, *J* = 15.6 Hz, 1H), 7.30 (s, 1H), 6.98 – 6.93 (m, 1H), 6.78 (s, 2H), 6.58 (s, 1H), 6.07 (dt, *J* = 9.6, 1.8 Hz, 1H), 4.21 – 4.16 (m, 2H), 4.04 (t, *J* = 6.5 Hz, 2H), 3.95 (s, 2H), 3.86 (s, 6H), 3.82 – 3.79 (m, 2H), 3.73 – 3.69 (m, 2H), 3.66 – 3.63 (m, 2H), 3.58 (t, *J* = 5.2 Hz, 2H), 3.51 – 3.46 (m, 2H), 3.01 (s, 2H), 2.93 – 2.88 (m, 2H), 2.51 – 2.46 (m, 2H), 2.39 – 2.34 (m, 1H), 2.22 – 2.16 (m, 2H), 1.91 – 1.86 (m, 4H), 1.25 (s, 9H). ^13^C NMR (151 MHz, CDCl_3_) δ 172.62, 170.32, 168.93, 166.07, 161.80, 158.84, 153.45, 145.68, 144.40, 143.66, 138.90, 130.77, 125.83, 121.24, 121.09, 120.16, 105.52, 72.29, 70.60, 70.38, 70.27, 70.05, 61.72, 56.21, 53.13, 42.05, 41.71, 38.79, 34.97, 31.44, 28.58, 28.50, 24.82. MS (ESI); m/z: [M+H]^+^ calcd for C_41_H_55_N_6_O_10_S_2_^+^: 855.3416, found 855.3371.

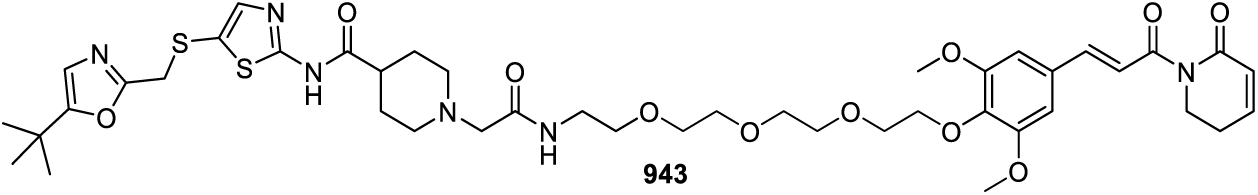

### (E)-N-(5-(((5-(tert-butyl)oxazol-2-yl)methyl)thio)thiazol-2-yl)-1-(14-(2,6- dimethoxy-4-(3-oxo-3-(6-oxo-3,6-dihydropyridin-1(2H)-yl)prop-1-en-1- yl)phenoxy)-2-oxo-6,9,12-trioxa-3-azatetradecyl)piperidine-4-carboxamide (943)

10.8 mg (0.012 mmol, 60% yield) was obtained as colorless gum from **8** (12 mg, 0.02 mmol) and **2** (9.9 mg, 0.02 mmol) by using the general procedure B. ^1^H NMR (600 MHz, Chloroform-*d*) δ 10.70 (br, 1H), 7.65 (d, *J* = 15.5 Hz, 1H), 7.51 – 7.46 (m, 1H), 7.40 (d, *J* = 15.5 Hz, 1H), 7.29 (s, 1H), 6.99 – 6.92 (m, 1H), 6.78 (s, 2H), 6.59 (s, 1H), 6.07 (dt, *J* = 9.7, 1.8 Hz, 1H), 4.20 – 4.16 (m, 2H), 4.04 (t, *J* = 6.5 Hz, 2H), 3.95 (s, 2H), 3.86 (s, 6H), 3.78 (t, *J* = 5.0 Hz, 2H), 3.73 – 3.61 (m, 8H), 3.57 (t, *J* = 5.1 Hz, 2H), 3.49 – 3.44 (m, 2H), 3.10 – 3.00 (m, 2H), 3.00 – 2.90 (m, 2H), 2.51 – 2.45 (m, 2H), 2.45 – 2.37 (m, 1H), 2.32 – 2.20 (m, 2H), 1.95 – 1.86 (m, 4H), 1.25 (s, 9H). ^13^C NMR (151 MHz, CDCl_3_) δ 172.68, 168.90, 166.00, 161.96, 161.78, 158.85, 153.42, 145.66, 144.27, 143.65, 138.79, 130.79, 125.81, 121.22, 120.93, 120.12, 105.49, 72.32, 70.51, 70.28, 70.05, 56.18, 53.10, 41.70, 38.78, 34.94, 31.44, 28.57, 24.81. MS (ESI); m/z: [M+H]^+^ calcd for C_43_H_59_N_6_O_11_S_2_ : 899.3678, found 899.3624.

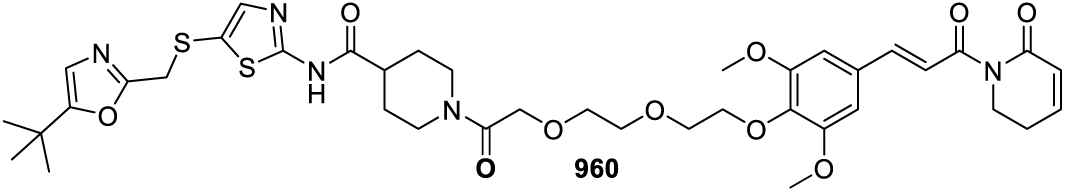

### (E)-N-(5-(((5-(tert-butyl)oxazol-2-yl)methyl)thio)thiazol-2-yl)-1-(2-(2-(2-(2,6- dimethoxy-4-(3-oxo-3-(6-oxo-3,6-dihydropyridin-1(2H)-yl)prop-1-en-1- yl)phenoxy)ethoxy)ethoxy)acetyl)piperidine-4-carboxamide (960)

12.0 mg (0.015 mmol, 46% yield) was obtained as colorless gum from **11** (16.0 mg, 0.032 mmol) and **1** (12.2 mg, 0.032 mmol) by using the general procedure B. ^1^H NMR (600 MHz, Chloroform-*d*) δ 10.46 (br, 1H), 7.65 (d, *J* = 15.5 Hz, 1H), 7.38 (d, *J* = 15.5 Hz, 1H), 7.30 (s, 1H), 6.99 – 6.93 (m, 1H), 6.78 (s, 2H), 6.58 (s, 1H), 6.13 (dt, *J* = 9.7, 1.8 Hz, 1H), 4.55 – 4.41 (m, 1H), 4.23 – 4.15 (m, 4H), 4.10 – 3.91 (m, 5H), 3.86 (s, 6H), 3.80 – 3.76 (m, 2H), 3.75 – 3.65 (m, 4H), 3.04 – 2.95 (m, 1H), 2.82 – 2.72 (m, 1H), 2.66 – 2.57 (m, 1H), 2.53 – 2.46 (m, 2H), 1.94 – 1.83 (m, 2H), 1.83 – 1.71 (m, 2H), 1.25 (s, 9H). ^13^C NMR (151 MHz, CDCl_3_) δ 171.96, 168.92, 167.79, 166.25, 161.79, 161.59, 158.82, 153.42, 145.80, 144.47, 143.67, 139.17, 130.54, 125.79, 121.19, 121.12, 120.13, 105.47, 72.39, 70.88, 70.69, 70.65, 70.50, 56.20, 44.06, 42.39, 41.75, 41.05, 34.96, 31.44, 28.57, 24.81. MS (ESI); m/z: [M+H]^+^ calcd for C_39_H_50_N_5_O_10_S_2_^+^: 812.2994, found 812.2946.

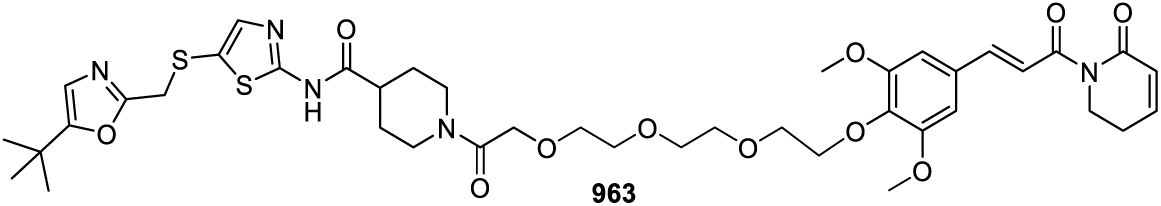

### (E)-N-(5-(((5-(tert-butyl)oxazol-2-yl)methyl)thio)thiazol-2-yl)-1-(2-(2-(2-(2-(2,6- dimethoxy-4-(3-oxo-3-(6-oxo-3,6-dihydropyridin-1(2H)-yl)prop-1-en-1- yl)phenoxy)ethoxy)ethoxy)ethoxy)acetyl)piperidine-4-carboxamide (963)

12.2 mg (0.014 mmol, 53% yield) was obtained as colorless gum from **12** (15.0 mg, 0.027 mmol) and **1** (10.3 mg, 0.027 mmol) by using the general procedure B.^1^H NMR (600 MHz, Chloroform-*d*) δ 10.41 (br, 1H), 7.61 (d, *J* = 15.5 Hz, 1H), 7.29 (s, 1H), 7.25 (d, *J* = 15.5 Hz, 1H), 6.98 (d, *J* = 9.7 Hz, 1H), 6.77 (s, 2H), 6.58 (s, 1H), 6.18 (d, *J* = 9.7 Hz, 1H), 4.25 – 4.13 (m, 4H), 4.06 – 3.97 (m, 2H), 3.93 (s, 2H), 3.88 (s, 6H), 3.81 – 3.67 (m, 11H), 3.62 – 3.55 (m, 1H), 3.07 – 2.99 (m, 1H), 2.67 – 2.59 (m, 2H), 2.54 – 2.48 (m, 2H), 1.90 – 1.75 (m, 4H), 1.25 (s, 9H). ^13^C NMR (151 MHz, CDCl_3_) δ 172.25, 169.06, 167.94, 166.99, 161.91, 161.55, 159.06, 153.50, 146.26, 145.08, 143.05, 137.28, 131.87, 125.82, 122.53, 120.82, 120.22, 105.07, 71.64, 69.87, 69.69, 69.62, 69.35, 69.17, 56.39, 43.03, 42.11, 41.29, 40.70, 35.11, 31.56, 28.69, 24.90. MS (ESI); m/z: [M+H]^+^ calcd for C_41_H_54_N_5_O_11_S_2_ : 856.3256, found 856.3203.

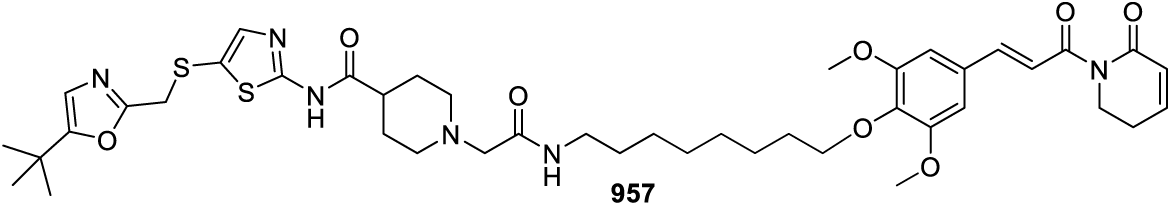

### (E)-N-(5-(((5-(tert-butyl)oxazol-2-yl)methyl)thio)thiazol-2-yl)-1-(2-((8-(2,6- dimethoxy-4-(3-oxo-3-(6-oxo-3,6-dihydropyridin-1(2H)-yl)prop-1-en-1- yl)phenoxy)octyl)amino)-2-oxoethyl)piperidine-4-carboxamide (957)

10.7 mg (0.013 mmol, 55% yield) was obtained as colorless gum from **9** (12.0 mg, 0.023 mmol) and **2** (11.4 mg, 0.023 mmol) by using the general procedure B. ^1^H NMR (600 MHz, Chloroform-*d*) δ 10.21 (br, 1H), 7.66 (d, *J* = 15.5 Hz, 1H), 7.40 (d, *J* = 15.5 Hz, 1H), 7.31 (s, 1H), 7.14 (s, 1H), 6.99 – 6.92 (m, 1H), 6.79 (d, *J* = 5.5 Hz, 2H), 6.59 (s, 1H), 6.06 (dt, *J* = 9.7, 1.8 Hz, 1H), 4.04 (t, *J* = 6.5 Hz, 2H), 3.99 (t, *J* = 6.7 Hz, 2H), 3.95 (s, 2H), 3.86 (s, 6H), 3.30 – 3.22 (m, 2H), 3.09 – 2.90 (m, 4H), 2.54 – 2.46 (m, 2H), 2.46 – 2.35 (m, 1H), 2.34 – 2.24 (m, 2H), 1.98 – 1.85 (m, 4H), 1.77 – 1.69 (m, 2H), 1.57 – 1.39 (m, 4H), 1.38 – 1.29 (m, 6H), 1.26 (s, 9H). ^13^C NMR (151 MHz, CDCl_3_) δ 168.97, 165.97, 161.82, 161.54, 158.82, 153.60, 145.59, 144.46, 143.92, 139.44, 130.36, 125.84, 121.27, 120.90, 120.13, 105.64, 73.58, 56.21, 53.14, 41.69, 39.03, 34.96, 31.45, 30.02, 29.63, 29.24, 29.18, 28.58, 26.87, 25.72, 24.82. MS (ESI); m/z: [M+H]^+^ calcd for C_43_H_59_N_6_O_8_S_2_^+^: 851.3830, found 851.3784.

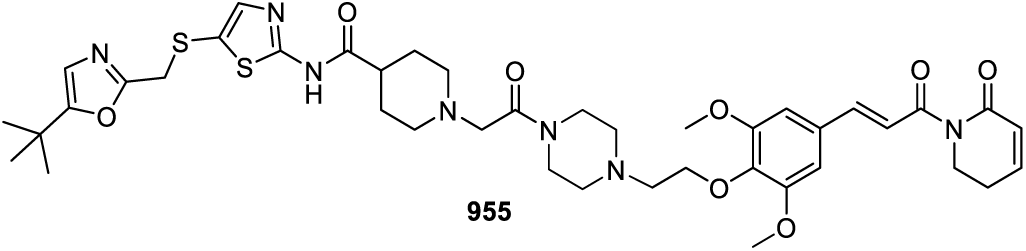

### (E)-N-(5-(((5-(tert-butyl)oxazol-2-yl)methyl)thio)thiazol-2-yl)-1-(2-(4-(2-(2,6- dimethoxy-4-(3-oxo-3-(6-oxo-3,6-dihydropyridin-1(2H)-yl)prop-1-en-1- yl)phenoxy)ethyl)piperazin-1-yl)-2-oxoethyl)piperidine-4-carboxamide (955)

57.3 mg (0.069 mmol, 49% yield) was obtained as white solid from **10** (74.0 mg, 0.14 mmol) and **2** (69.2 mg, 0.14 mmol) by using the general procedure B.^1^H NMR (600 MHz, Methanol-*d*_4_) δ 7.61 (d, *J* = 15.6 Hz, 1H), 7.38 (d, *J* = 15.6 Hz, 1H), 7.31 (s, 1H), 7.11-7.04 (m, 1H), 6.92 (s, 2H), 6.67 (s, 1H), 6.04 – 5.98 (m, 1H), 4.16 – 4.09 (m, 2H), 4.02 – 3.94 (m, 4H), 3.88 (s, 6H), 3.70 – 3.65 (m, 2H), 3.65 – 3.58 (m, 2H), 3.24 (s, 2H), 3.02 – 2.92 (m, 2H), 2.81 (t, *J* = 5.3 Hz, 2H), 2.72 – 2.66 (m, 2H), 2.66 – 2.59 (m, 2H), 2.55 – 2.48 (m, 2H), 2.47 – 2.42 (m, 1H), 2.21 – 2.11 (m, 2H), 1.87 – 1.77 (m, 4H), 1.23 (s, 9H). ^13^C NMR (151 MHz, MeOD) δ 175.24, 170.56, 167.76, 163.46, 163.31, 161.24, 154.97, 148.41, 146.56, 144.18, 139.99, 132.48, 125.89, 122.73, 121.07, 120.68, 106.56, 71.05, 61.22, 58.70, 56.66, 53.86, 46.29, 43.03, 42.66, 35.17, 32.44, 28.90, 25.75. MS (ESI); m/z: [M+H]^+^ calcd for C_41_H_54_N7O_8_S_2_^+^: 836.3470, found 836.3424.

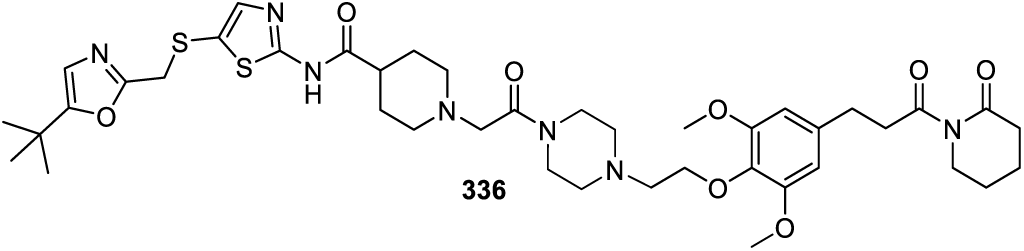

### N-(5-(((5-(tert-butyl)oxazol-2-yl)methyl)thio)thiazol-2-yl)-1-(2-(4-(2-(2,6- dimethoxy-4-(3-oxo-3-(2-oxopiperidin-1-yl)propyl)phenoxy)ethyl)piperazin-1-yl)- 2-oxoethyl)piperidine-4-carboxamide (336)

8.3 mg (0.001 mmol, 52% yield) was obtained as colorless gum from **13** (10.0 mg, 0.019 mmol) and **2** (9.4 mg, 0.019 mmol) by using the general procedure B. ^1^H NMR (600 MHz, Chloroform-*d*) δ 10.51 (br, 1H), 7.31 (s, 1H), 6.60 (s, 1H), 6.44 (s, 2H), 4.07 (t, *J* = 5.6 Hz, 2H), 3.96 (s, 2H), 3.82 (s, 6H), 3.74 – 3.64 (m, 6H), 3.30 (s, 2H), 3.24 – 3.20 (m, 2H), 3.04 (d, *J* = 11.3 Hz, 2H), 2.90 (t, *J* = 7.7 Hz, 2H), 2.80 (t, *J* = 5.5 Hz, 2H), 2.62 (dt, *J* = 22.2, 5.1 Hz, 4H), 2.57 – 2.52 (m, 2H), 2.42 (s, 1H), 2.28 (s, 2H), 1.95 – 1.80 (m, 8H), 1.25 (s, 9H). ^13^C NMR (151 MHz, CDCl_3_) δ 176.16, 173.54, 172.63, 167.45, 161.83, 161.79, 158.94, 153.17, 144.40, 137.26, 135.04, 121.12, 120.12, 105.44, 69.95, 61.08, 57.87, 56.05, 52.75, 45.37, 44.10, 43.69, 41.39, 34.94, 34.91, 31.56, 31.45, 28.57, 22.44, 20.27. MS (ESI); m/z: [M+H]^+^ calcd for C_41_H_58_N_7_O_8_S_2_ : 840.3783, found 840.3732.

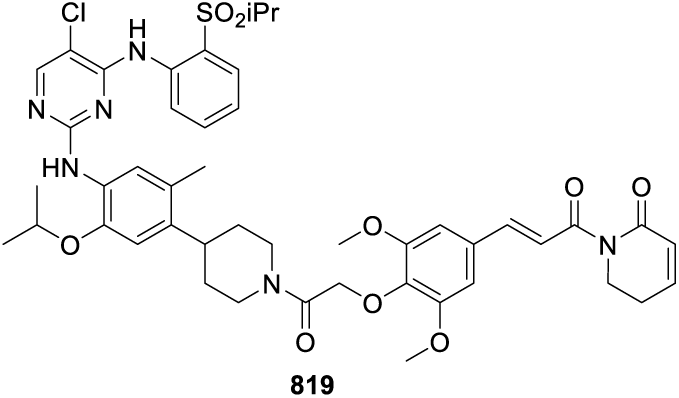

### (E)-1-(3-(4-(2-(4-(4-((5-chloro-4-((2-(isopropylsulfonyl)phenyl)amino)pyrimidin- 2-yl)amino)-5-isopropoxy-2-methylphenyl)piperidin-1-yl)-2-oxoethoxy)-3,5- dimethoxyphenyl)acryloyl)-5,6-dihydropyridin-2(1H)-one (819)

4.3 mg (0.005 mmol, 27% yield) was obtained as colorless gum from **14** (7.6 mg, 0.018 mmol) and Ceritinib (10.2 mg, 0.018 mmol) by using the general procedure B. ^1^H NMR (600 MHz, CDCl3) δ 9.50 (s, 1H), 8.58 (d, *J* = 8.3 Hz, 1H), 8.16 (s, 1H), 8.03 (s, 1H), 7.94 (dd, *J* = 8.0, 1.5 Hz, 1H), 7.67 (d, *J* = 15.5 Hz, 1H), 7.65 – 7.60 (m, 1H), 7.55 (s, 1H), 7.43 (d, *J* = 15.5 Hz, 1H), 7.29 – 7.24 (m, 1H), 6.98 – 6.92 (m, 1H), 6.80 (s, 2H), 6.72 (s, 1H), 6.05 (dt, *J* = 9.7, 1.7 Hz, 1H), 4.80 (d, *J* = 13.3 Hz, 1H), 4.72 (d, *J* = 12.0 Hz, 1H), 4.65 (d, *J* = 12.0 Hz, 1H), 4.58 – 4.50 (m, 2H), 4.05 (t, *J* = 6.5 Hz, 2H), 3.88 (s, 6H), 3.27 (dt, *J* = 13.7, 6.9 Hz, 1H), 3.21 (t, *J* = 12.5 Hz, 1H), 2.98 – 2.92 (m, 1H), 2.73 (t, *J* = 12.1 Hz, 1H), 2.51 – 2.46 (m, 2H), 2.20 (s, 3H), 1.84 (t, *J* = 9.5 Hz, 2H), 1.74 – 1.62 (m, 2H), 1.36 (d, *J* = 6.0 Hz, 6H), 1.32 (d, *J* = 6.9 Hz, 6H). ^13^C NMR (151 MHz, CDCl_3_) δ 168.82, 166.51, 165.88, 157.46, 155.38, 155.34, 153.34, 145.60, 144.73, 143.55, 138.50, 138.01, 136.79, 134.64, 131.41, 131.29, 127.83, 126.84, 125.82, 124.93, 123.68, 123.12, 121.47, 120.67, 110.83, 105.84, 105.43, 72.11, 71.63, 56.21, 55.46, 46.38, 43.06, 41.66, 38.40, 33.46, 32.32, 24.81, 22.32, 22.22, 19.02, 15.38. MS (ESI); m/z: [M+H]^+^ calcd for C_46_H_54_ClN_6_O_9_S_2_^+^: 901.3356, found 901.3356.

### NMR and HRMS spectra for 955, 336 and 819

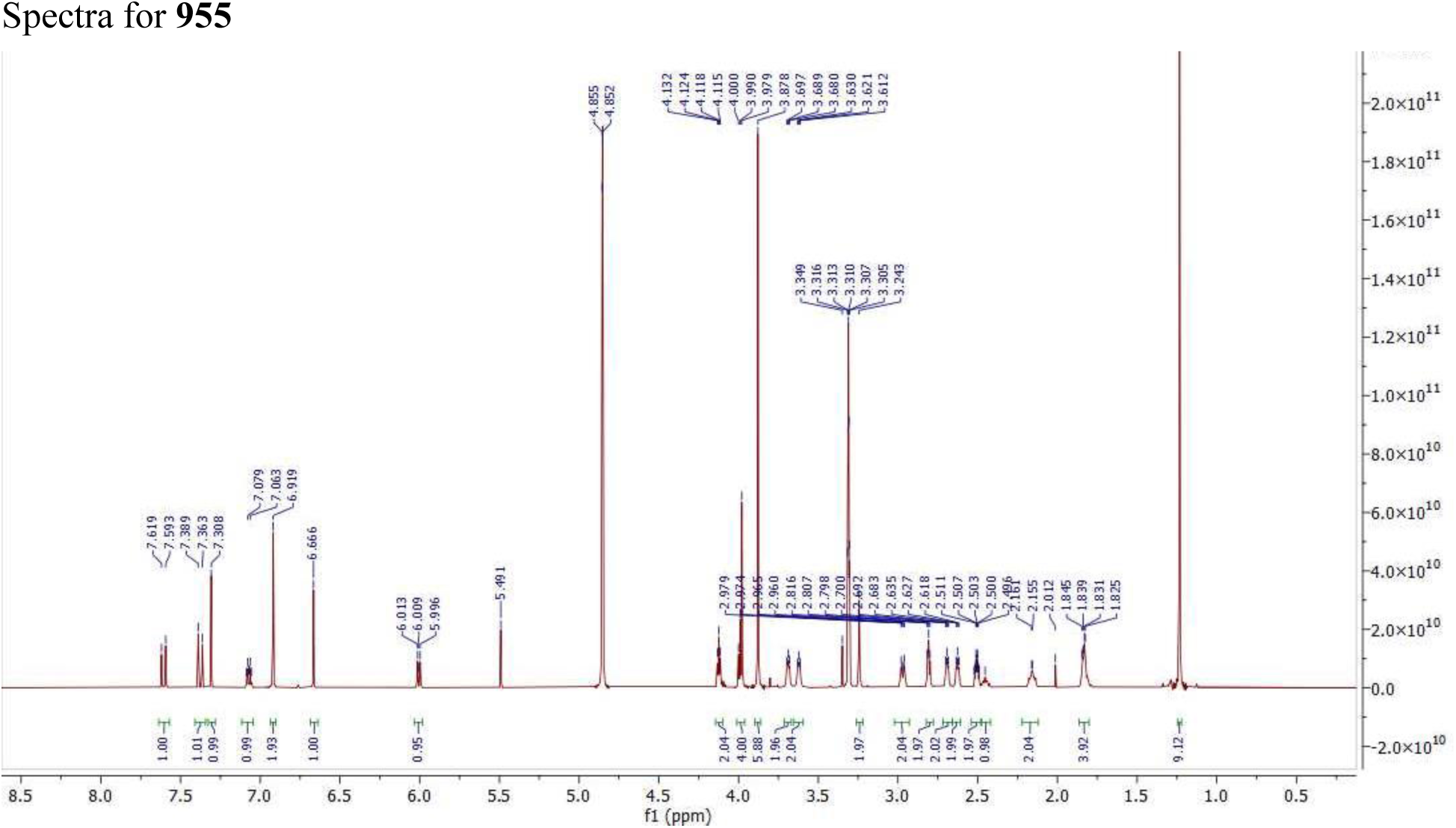
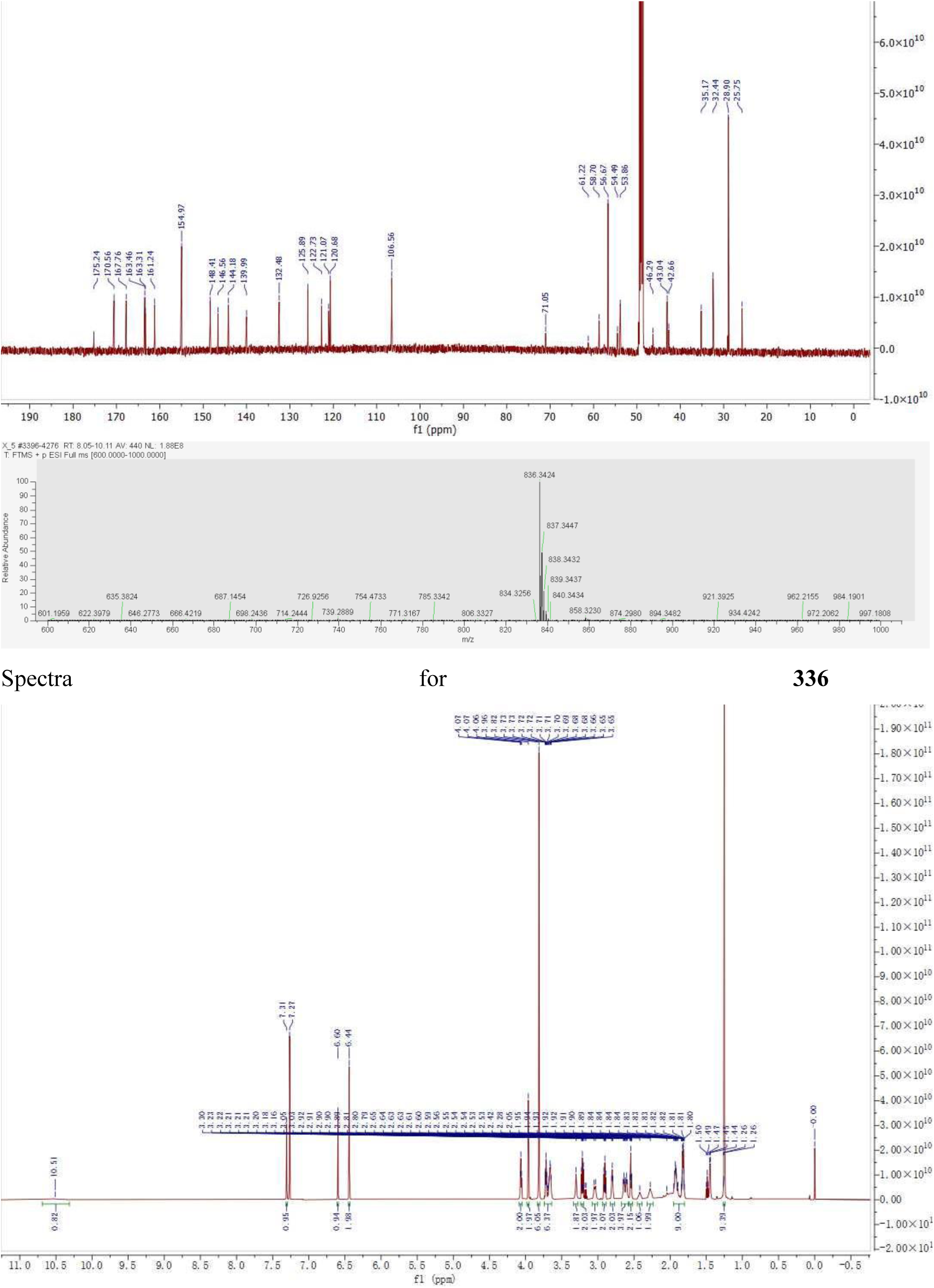
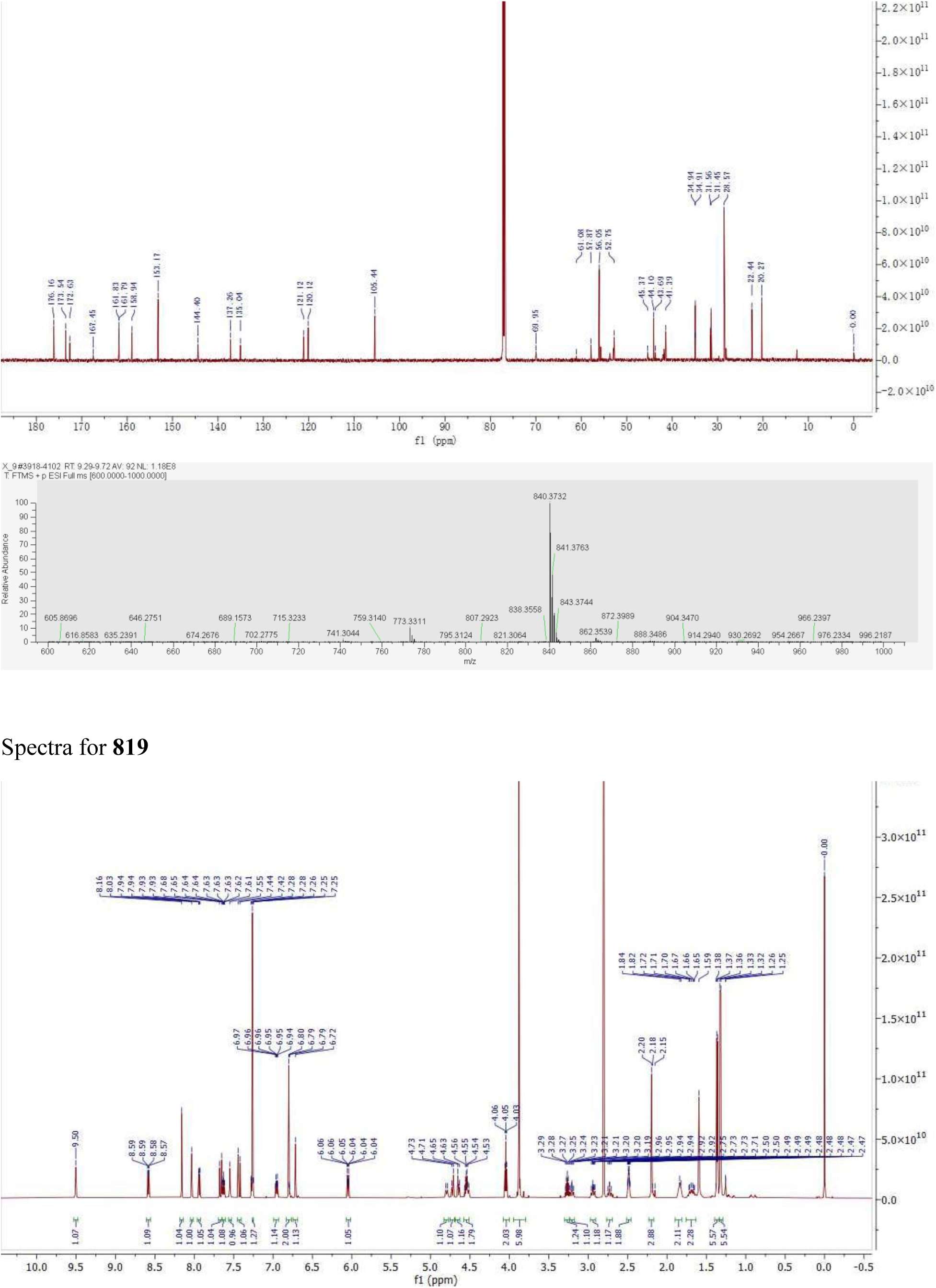
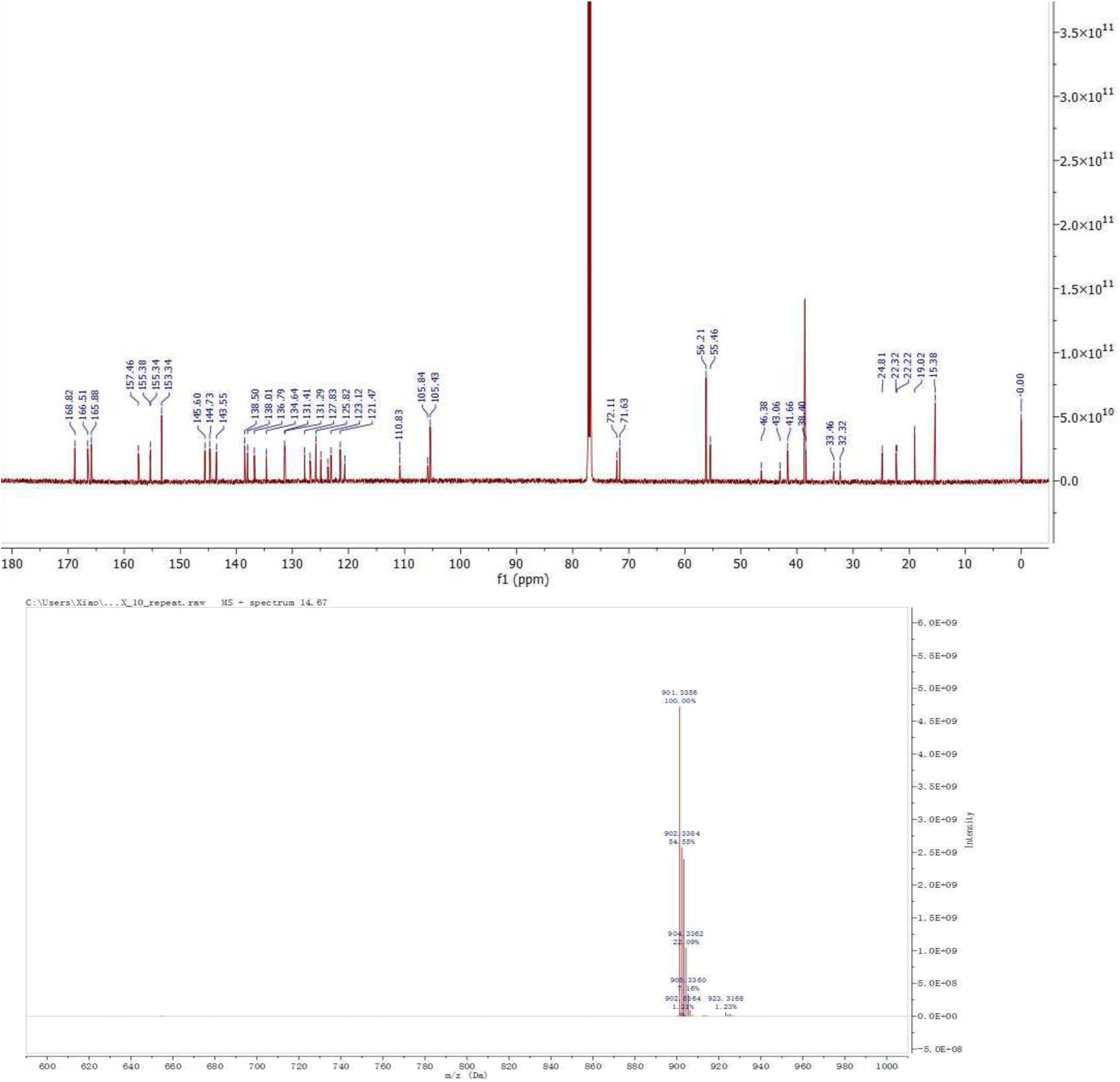

## QUANTIFICATION AND STATISTICAL ANALYSIS

Immunoblot data were quantified using the ImageJ (v1.53a) software from NIH. Data analysis was performed using GraphPad Prism software (version 9) unless indicated otherwise. The n number for each experiment is indicated in the figure legends. All graphs presented represent the mean ± standard deviation of the mean (SD) unless otherwise stated.

