## Supplemental Figures for "Piperlongumine (PL) conjugates induce targeted protein degradation"

[†] These authors contributed equally.

**LEAD CONTACT**

**Supplementary Figures**

**Figure S1. Validation of PL-binding E3 ligases (related to Table S1 and STAR Methods).** (A) The structures of PL and PL-Alkyne probe; (B) The workflow of the competitive ABPP assay; (C) Western blot to detect biotin-labelled proteins in the streptavidin pull-downed samples; and (D) The PL-binding proteins identified by mass spectrometry. E3 ligases are highlighted in red and GSTO1 is shown in green. MOLT4 is a human T-cell acute lymphoblastic leukemia (T-ALL) cell line. (E-H) GO and KEGG enrichment analysis of PL binding proteins. (E) Enriched GO BP items; (F) Enriched GO MF items; (G) Enriched GO CC items; (H) Enriched KEGG items. The GO items or KEGG items with FDR<0.05 were extracted and shown.

**Figure S2. Synthesis and evaluation of a series of PL-SNS-032 conjugates (related to Figure 1, Chemistry Methods and STAR Methods).** (A) The general chemical structure of PL-SNS-032 conjugates; (B) Evaluation of the biological activites of different PL-SNS-032 conjugates in MOLT4 cells. The $DC_{50}$ and $D_{max}$ were calculated using Prism based on the ImageJ-quantified western blots in panel C and the $EC_{50}$ were determined by MTS assays. For the MTS assays, the data are presented as mean ± SD from three independent experiments; (C) The CDK9 degradation profiling of the PL-SNS-032 conjugates evaluated in MOLT4 cells. Representative immunoblots are shown and β-actin was used as a loading control in all immunoblot analyses. The quantification of the relative CDK9 protein content in the immunoblots is presented as mean ± SD ($n = 2$ biologically independent experiments) in the bar graph (bottom panel). Statistical significance was calculated with unpaired two-tailed Student's *t*-tests. $^{*}P < 0.05$; $^{**}P < 0.01$.

**Figure S3. 955 degrades CDK9 in 293T and K562 cells in a time- and UPS- dependent manner (related to Figure 1, Figure 2 and STAR Methods).** (A) Time course of **955**-induced CDK9 degradation in 293T cells. (B) Time course of **955**-induced CDK9 degradation in K562 cells. Pretreatment with proteasome inhibitor (MG132 or Bortezomib) blocks CDK9 degradation by **955** in 293T (C) and K562 (E) cells. Pretreatment with neddylation inhibitor MLN4924 blocks CDK9 degradation by **955** in 293T (D) and K562 (F) cells. Representative immunoblots are shown and β-actin was used as a loading control in all immunoblot analyses. The quantification of the relative CDK9 protein content in the immunoblots is presented as mean ± SD ($n = 2$ biologically independent experiments) in the bar graph (bottom panel).

Statistical significance was calculated with unpaired two-tailed Student’s *t*-tests. *$P < 0.05$; **$P < 0.01$; NS: not significant.

**Figure S4. KEAP1 knockdown blocks 955-induced CDK9 degradation (related to Figure 3 and STAR Methods).** KEAP1 (A), TRIP12 (B), or TRAF6 (C) was knocked down by siRNAs in H1299 cells and then the cells were either untreated or treated with indicated concentration of **955** for 6 h. (D) KEAP1 single or KEAP1 and NRF2 double knockdown by siRNAs in H1299 cells and then the cells were either untreated or treated with indicated concentration of **955** for 6 h. Representative immunoblots are shown and β-actin was used as a loading control in all immunoblot analyses. The quantification of the relative CDK9 protein content in the immunoblots is presented as mean ± SD ($n = 2$ biologically independent experiments) in the bar graph (bottom panel). Statistical significance was calculated with unpaired two-tailed Student’s *t*-tests. *$P < 0.05$; **$P < 0.01$; NS: not significant.

**Figure S5. Analysis of TMT-proteomics and RNA-seq data (related to Figure 4, Figure 5, Table S3 and Table S4).** (A) Venn diagram to show the differentially expressed proteins in MOLT4 cells after 6 h of treatment with 955 and SNS-032. (B-C) GO analysis of the transcriptional changes in MOLT4 cells treated with 0.1 μM 955 or 1 μM SNS-032 for 6 h. GO enrichment analysis of down- (B) and up- (C) regulated differentially expressed genes identified from 955 and SNS-032 treatments. GO BP items with FDR<0.05 were extracted and shown.

**Figure S6. Percentages of cell killing in LNCaP and H1299 cells with or without KEAP1 kncokdown (related to Figure 5 and STAR Methods).** LNCaP and H1299 cells were treated with control siRNA or siRNAs targeting KEAP1 for 48 h and then reseeded and treated with indicated compound (0.1 μM for LNCaP cells and 0.3 μM for H1299 cells) for 24 h. The cell viability was then determined by MTS assay. The data are presented as mean ± SD from three replicate cell cultures in a representative experiment. Data are representative of two independent experiments. Statistical significance was calculated with unpaired two-tailed Student’s *t*-tests. **$P < 0.01$; ***$P < 0.001$; NS: not significant.

### Supplement Figures

#### Figure S1

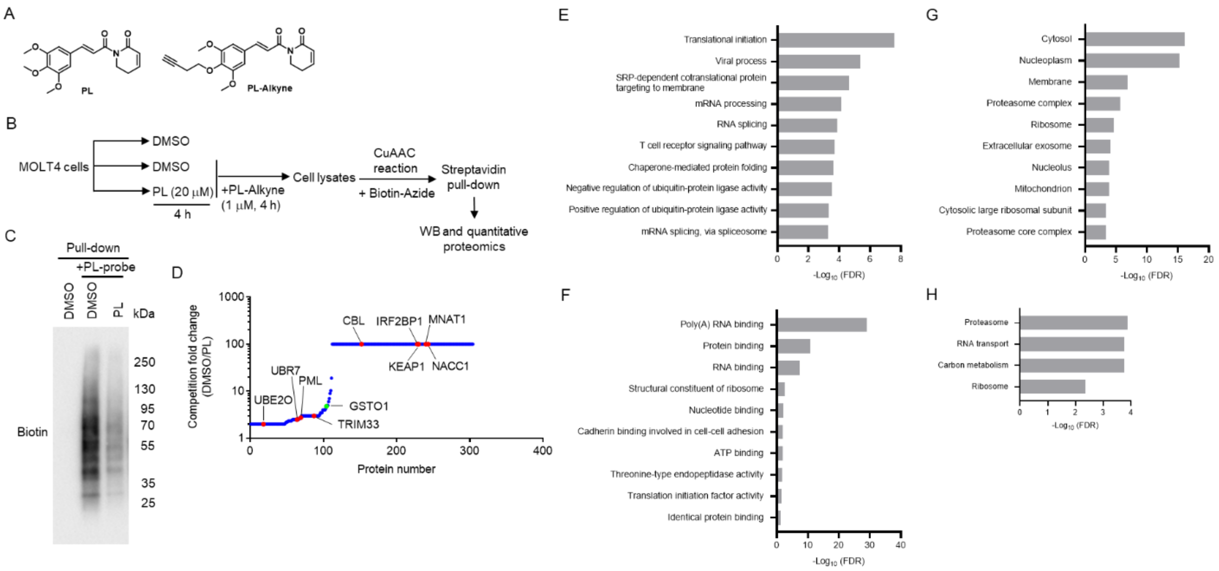

### Figure S2

A

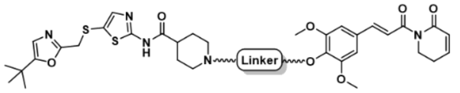

B

| Compound | Linker | $DC_{50}$ (nM) | $D_{max}$ | $EC_{50}$ (nM) |
|---|---|---|---|---|
| 941 | 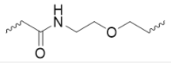 | 15 | 87 | 120 ± 9 |
| 929 | 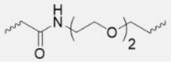 | 12 | 93 | 138 ± 10 |
| 943 | 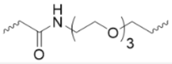 | 10 | 96 | 132 ± 1 |
| 957 | 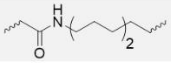 | 102 | 74 | 318 ± 15 |
| 960 | 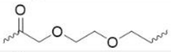 | 28 | 87 | 119 ± 7 |
| 963 | 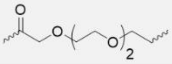 | 15 | 92 | 107 ± 8 |
| 955 | 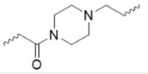 | 9 | 96 | 28 ± 5 |
| PL | | NA | NA | >3000 |
| SNS-032 | | NA | NA | 197 ± 8 |
| PL+SNS032 | | NA | NA | 174 ± 38 |

C

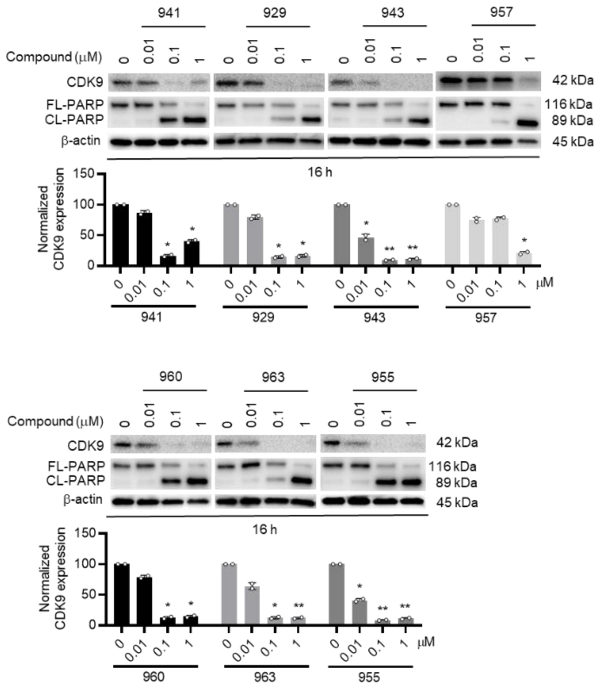

**Figure S3**

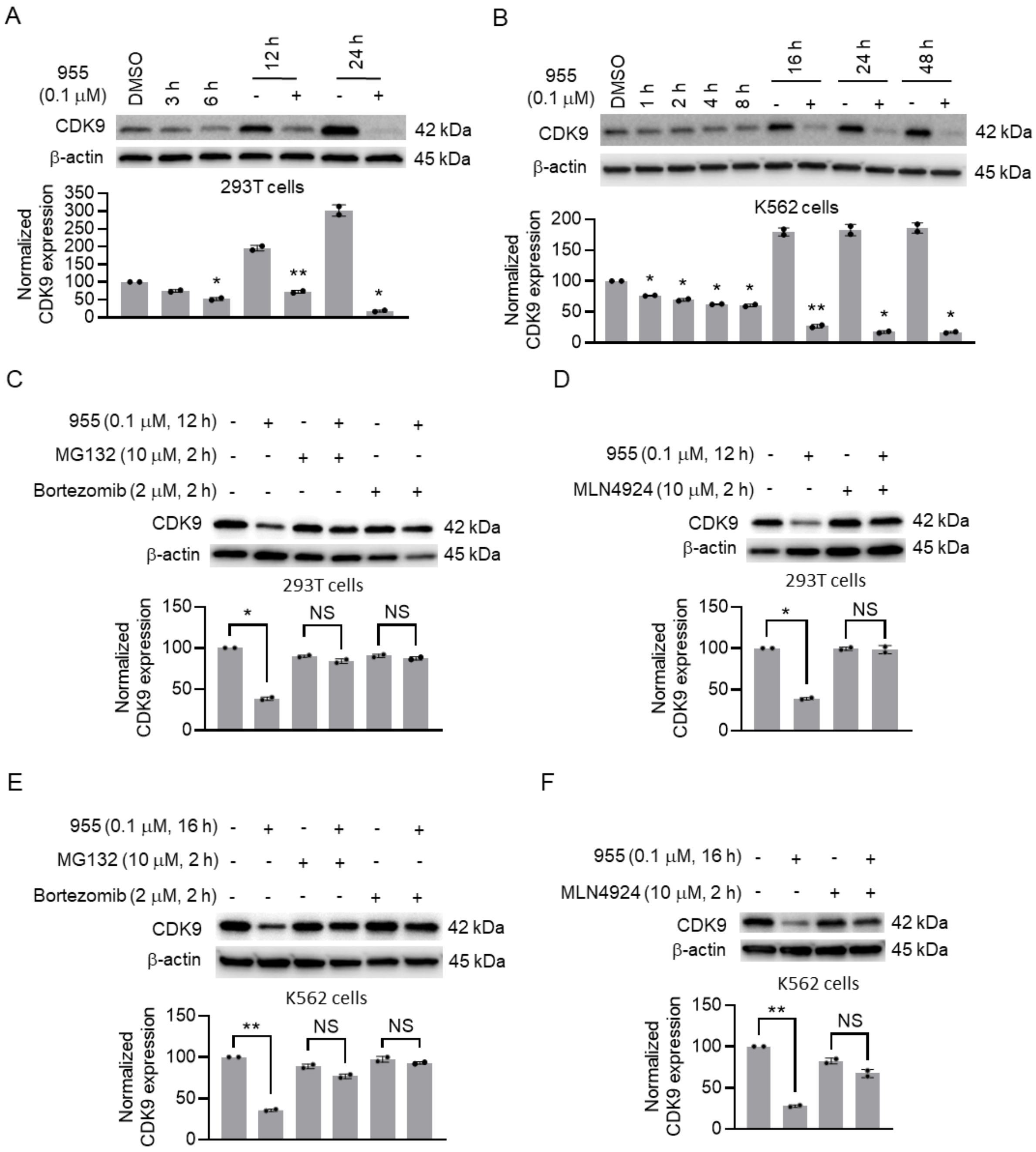

A
955 (0.1 μM)
DMSO
3 h
6 h
12 h
24 h
CDK9
β-actin
42 kDa
45 kDa
293T cells
Normalized CDK9 expression
B
955 (0.1 μM)
DMSO
1 h
2 h
4 h
8 h
16 h
24 h
48 h
CDK9
β-actin
42 kDa
45 kDa
K562 cells
Normalized CDK9 expression
C
955 (0.1 μM, 12 h)
MG132 (10 μM, 2 h)
Bortezomib (2 μM, 2 h)
CDK9
β-actin
42 kDa
45 kDa
293T cells
NS
Normalized CDK9 expression
D
955 (0.1 μM, 12 h)
MLN4924 (10 μM, 2 h)
CDK9
β-actin
42 kDa
45 kDa
293T cells
NS
Normalized CDK9 expression
E
955 (0.1 μM, 16 h)
MG132 (10 μM, 2 h)
Bortezomib (2 μM, 2 h)
CDK9
β-actin
42 kDa
45 kDa
K562 cells
NS
Normalized CDK9 expression
F
955 (0.1 μM, 16 h)
MLN4924 (10 μM, 2 h)
CDK9
β-actin
42 kDa
45 kDa
K562 cells
NS
Normalized CDK9 expression

**Figure S4**

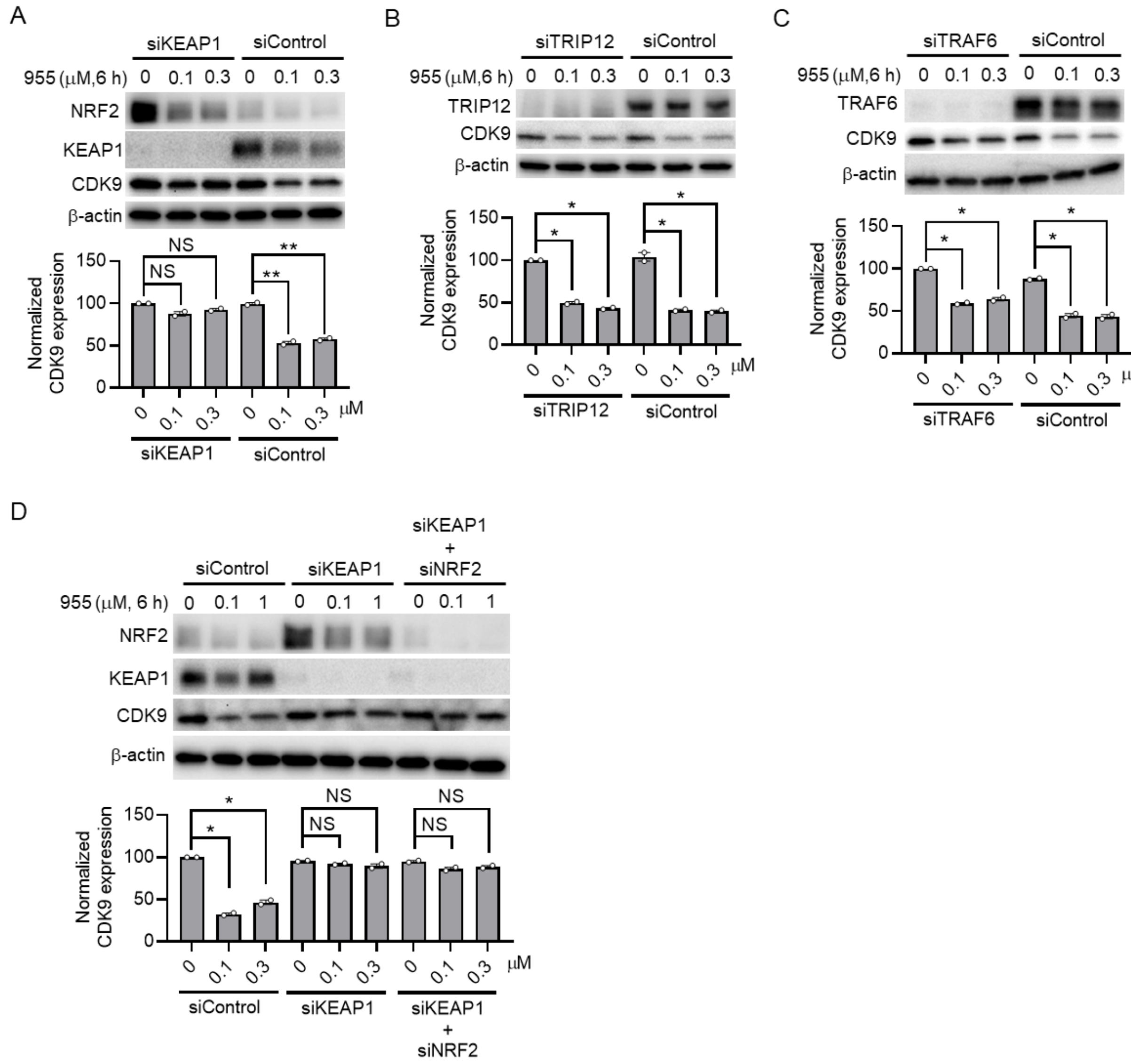
A
siKEAP1
siControl
955 (μM,6 h) 0 0.1 0.3 0 0.1 0.3
NRF2
KEAP1
CDK9
β-actin
Normalized CDK9 expression
150
100
50
0
NS
NS
**
**
0 0.1 0.3 0 0.1 0.3 μM
siKEAP1
siControl
B
siTRIP12
siControl
955 (μM,6 h) 0 0.1 0.3 0 0.1 0.3
TRIP12
CDK9
β-actin
Normalized CDK9 expression
150
100
50
0
*
*
*
*
0 0.1 0.3 0 0.1 0.3 μM
siTRIP12
siControl
C
siTRAF6
siControl
955 (μM,6 h) 0 0.1 0.3 0 0.1 0.3
TRAF6
CDK9
β-actin
Normalized CDK9 expression
150
100
50
0
*
*
*
*
0 0.1 0.3 0 0.1 0.3 μM
siTRAF6
siControl
D
siControl
siKEAP1
siKEAP1 + siNRF2
955 (μM, 6 h) 0 0.1 1 0 0.1 1 0 0.1 1
NRF2
KEAP1
CDK9
β-actin
Normalized CDK9 expression
150
100
50
0
*
*
NS
NS
NS
NS
0 0.1 0.3 0 0.1 0.3 0 0.1 0.3 μM
siControl
siKEAP1
siKEAP1 + siNRF2

**Figure S5**

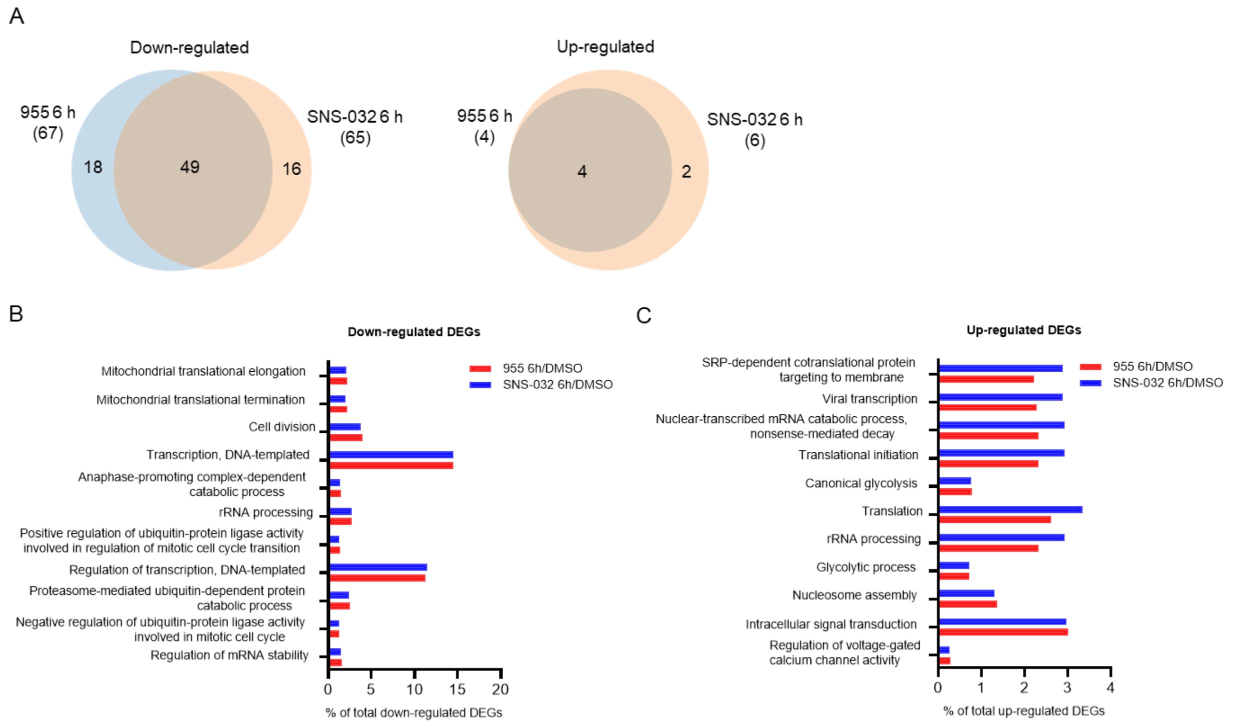
A
Down-regulated
955 6 h
(67)
SNS-032 6 h
(65)
18
49
16
Up-regulated
955 6 h
(4)
SNS-032 6 h
(6)
4
2
B
Down-regulated DEGs
Mitochondrial translational elongation
Mitochondrial translational termination
Cell division
Transcription, DNA-templated
Anaphase-promoting complex-dependent catabolic process
rRNA processing
Positive regulation of ubiquitin-protein ligase activity involved in regulation of mitotic cell cycle transition
Regulation of transcription, DNA-templated
Proteasome-mediated ubiquitin-dependent protein catabolic process
Negative regulation of ubiquitin-protein ligase activity involved in mitotic cell cycle
Regulation of mRNA stability
955 6h/DMSO
SNS-032 6h/DMSO
0 5 10 15 20
% of total down-regulated DEGs
C
Up-regulated DEGs
SRP-dependent cotranslational protein targeting to membrane
Viral transcription
Nuclear-transcribed mRNA catabolic process, nonsense-mediated decay
Translational initiation
Canonical glycolysis
Translation
rRNA processing
Glycolytic process
Nucleosome assembly
Intracellular signal transduction
Regulation of voltage-gated calcium channel activity
955 6h/DMSO
SNS-032 6h/DMSO
0 1 2 3 4
% of total up-regulated DEGs

**Figure S6**

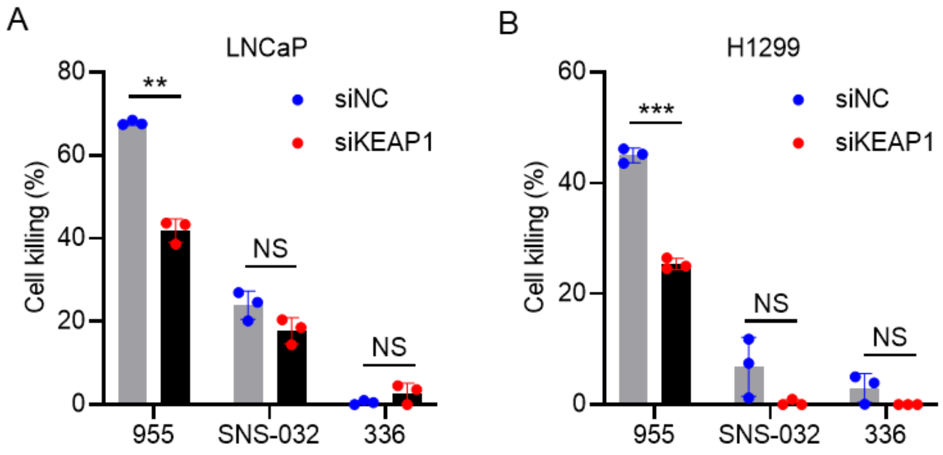

A
LNCaP
80
60
40
20
0
Cell killing (%)
**
NS
NS
siNC
siKEAP1
955
SNS-032
336
B
H1299
60
40
20
0
Cell killing (%)
***
NS
NS
siNC
siKEAP1
955
SNS-032
336
